# Altered Metabolism and DAM-signatures in Female Brains and Microglia with Aging

**DOI:** 10.1101/2023.11.28.569104

**Authors:** Nicholas R W Cleland, Garrett J Potter, Courtney Buck, Daphne Quang, Dean Oldham, Mikaela Neal, Anthony Saviola, Christy S. Niemeyer, Evgenia Dobrinskikh, Kimberley D. Bruce

## Abstract

Despite Alzheimer’s disease (AD) disproportionately affecting women, the mechanisms remain elusive. In AD, microglia undergo ‘metabolic reprogramming’, which contributes to microglial dysfunction and AD pathology. However, how sex and age contribute to metabolic reprogramming in microglia is understudied. Here, we use metabolic imaging, transcriptomics, and metabolic assays to probe age-and sex-associated changes in brain and microglial metabolism. Glycolytic and oxidative metabolism in the whole brain was determined using Fluorescence Lifetime Imaging Microscopy (FLIM). Young female brains appeared less glycolytic than male brains, but with aging, the female brain became ‘male-like.’ Transcriptomic analysis revealed increased expression of disease-associated microglia (DAM) genes (e.g., *ApoE*, *Trem2*, *LPL*), and genes involved in glycolysis and oxidative metabolism in microglia from aged females compared to males. To determine whether estrogen can alter the expression of these genes, BV-2 microglia-like cell lines, which abundantly express DAM genes, were supplemented with 17β-estradiol (E2). E2 supplementation resulted in reduced expression of DAM genes, reduced lipid and cholesterol transport, and substrate-dependent changes in glycolysis and oxidative metabolism. Consistent with the notion that E2 may suppress DAM-associated factors, LPL activity was elevated in the brains of aged female mice. Similarly, DAM gene and protein expression was higher in monocyte-derived microglia-like (MDMi) cells derived from middle-aged females compared to age-matched males and was responsive to E2 supplementation. FLIM analysis of MDMi from young and middle-aged females revealed reduced oxidative metabolism and FAD+ with age. Overall, our findings show that altered metabolism defines age-associated changes in female microglia and suggest that estrogen may inhibit the expression and activity of DAM-associated factors, which may contribute to increased AD risk, especially in post-menopausal women.

## 1. Introduction

Alzheimer’s Disease (AD) is the most common cause of dementia in the United States, affecting 73% of Americans aged 75 or over (Rajan et al., 2021). AD neuropathogenesis involves the accumulation of amyloid-beta (Aβ) plaques, neurofibrillary tangles (NFTs) (Wood et al., 1986), neuroinflammation (Calsolaro and Edison, 2016), glial cell dysfunction (De Sousa, 2022), lipid droplet (LD) accumulation (Marschallinger et al., 2020), aberrant lipid and lipoprotein processing (Cleland et al., 2021; Kao et al., 2020), and brain glucose hypometabolism (Abeysinghe et al., 2020; Johri, 2021; Mullins et al., 2018) leading to neurodegeneration (Masters et al., 1985; Serrano-Pozo et al., 2011; Wood et al., 1986) and cognitive decline (Porsteinsson et al., 2021). Despite the complexity of AD, current FDA-approved therapies only target one, if any, of the underlying facets of the disease and may not be available to, or effective, in many subjects with AD (Liu and Howard, 2021; Manly and Glymour, 2021; Osborne et al., 2023; Ramanan and Day, 2023; van Dyck et al., 2023). Hence, there is a clear need to identify alternative strategies that can simultaneously target many of the underlying mechanisms contributing to AD pathology and may improve outcomes for those living with or at risk of developing AD.

Microglia—the key immune cells of the brain—regulate many of the processes leading to AD pathology and disease onset. For example, microglia phagocytose amyloid beta (Aβ) (Grubman et al., 2021), produce inflammatory cytokines (Heneka et al., 2015), accumulate LDs (Marschallinger et al., 2020), regulate lipid and lipoprotein metabolism (Cleland et al., 2021), and contribute to neuronal loss and regeneration (Lloyd et al., 2019). Hence, microglia have the capacity to protect against or promote the development of AD depending on their activation state and function and are an emerging target in the search for novel AD therapeutics. It is becoming increasingly recognized that microglial activation is intrinsically linked to metabolic status (Fumagalli et al., 2018; Lynch, 2020; Minhas et al., 2021). When microglia become activated, they shift towards glucose utilization and away from oxidative phosphorylation for ATP production (Cleland et al., 2021; Lynch, 2020). Chronic activation eventually leads to “metabolic reprogramming,” when glycolytic shifts cannot sustain the bioenergetic needs of the cell, contributing to AD pathology by impairing the cell’s capacity to clear Aβ, increasing the production of inflammatory cytokines and promoting LD accumulation and neurodegeneration (Baik et al., 2019; Preeti et al., 2022; Shippy and Ulland, 2020). While there is growing support for the immunometabolic hypothesis of microglial dysfunction in AD, what triggers the metabolic reprogramming of microglia in the first place remains understudied (Chen et al., 2023; Fairley et al., 2021; Jung et al., 2022).

Two-thirds of Americans living with AD are female (Rajan et al., 2021). While younger females appear to be somewhat protected, the lifetime risk of developing AD is almost double in females over the age of 45, highlighting the impact of age on disease onset (Rajan et al., 2021). Emerging studies suggest that sex differences in microglial metabolism play a role in the increased incidence of AD in females (Demarest et al., 2020; Kodama and Gan, 2019; Park et al., 2023; Zhao et al., 2016). Throughout life, microglia exhibit sex- and age-specific differences in processes key to microglial function such as chemotaxis and cell migration (Thion et al., 2018; Villa et al., 2018), the response to inflammatory stimuli (Lynch, 2022; Nelson et al., 2017; Schwarz et al., 2012; Thion et al., 2018; Yanguas-Casás et al., 2018), and phagocytosis (Hammond et al., 2019; Hanamsagar et al., 2017; Loram et al., 2012; Lynch, 2022; Schwarz et al., 2012). In rodent models of AD, female microglia are more glycolytic, and less phagocytic than male mice (Guillot-Sestier et al., 2021). In addition, recent studies have shown sex-specific changes in senescence and disease associated microglia (DAM) signatures, which become more pronounced with aging (Ocanas et al., 2023). Therefore, microglia in aged females exhibit exacerbated metabolic reprogramming associated with AD pathology. However, why female microglia show exacerbated metabolic changes with aging and disease, and whether this translates to functional metabolic changes remains unknown.

Microglia are highly responsive to their environment and are profoundly affected by hormones such as 17β-estradiol (E2) (Vegeto et al., 2001). Indeed, the local production of E2 by astrocytes is one of the fist responses of both male and female brain tissue to injury (Duncan and Saldanha, 2011). A number of studies have shown the suppressive effect of E2 on microglial activation (Loiola et al., 2019; Slowik et al., 2018; Vegeto et al., 2000; Yun et al., 2018), production of reactive oxygen species (Bruce-Keller et al., 2000), and inflammatory mediators (Dimayuga et al., 2005; Wang et al., 2021). Given that there is a temporal relationship between menopause and the onset of AD, and sexually dimorphic microglial dysfunction is more pronounced with aging, it is likely that falling E2 concentrations with aging may alter microglial metabolism and function to exacerbate AD risk. In line with this, the use of hormone replacement therapy (HRT) during and after menopause is being considered as viable strategy to reduce AD risk. However, results of clinical trials have yielded inconsistent results (Grady et al., 2002; Mulnard et al., 2000). Therefore, empirical studies defining the effect of E2 on microglial metabolism are much needed.

Here we sought to investigate age-and sex-associated changes in brain and microglial metabolism that may contribute to AD risk in women. Glycolysis and oxidative metabolism in the whole brain was determined using Fluorescence Lifetime Imaging Microscopy (FLIM). Interestingly, female brains showed a more pronounced increase in glycolysis with aging than male brains. Microglia from aged female brains showed increased expression of disease-associated microglia (DAM) genes, and genes involved in glycolysis and oxidative metabolism. E2 supplementation in microglial cell lines resulted in reduced expression of DAM genes, intracellular retention of LPL, reduced cholesterol efflux, and substrate-dependent changes in glycolysis and oxidative metabolism. These data suggest that E2 may have an inhibitory effect on the expression and activity of DAM-associated factors. Consistent with this notion, we also observed elevated LPL activity in the brains of aged female mice compared to aged males. Studies using human monocyte-derived microglia-like (MDMi) cells revealed higher DAM and GLUT5 gene expression in cells derived from middle-aged females compared to males, which could be partially rescued by E2 supplementation. Using FLIM as a functional measure of metabolism in MDMi, we revealed metabolic changes associated with aging in human microglia. Our findings support the notion that altered metabolism defines age-associated changes in female microglia, and supports the need for further studies aimed at restoring microglial metabolism to improve outcomes for women with, or at risk of, AD.

## 2. Results

### 2.1 Increased glycolysis and reduced mitochondrial function in aged female brains

We asked whether there were differences in brain metabolism between males and females, and whether these changes were exacerbated with aging. Many studies investigating brain metabolism have extrapolated their findings from transcriptomic analyses, which is suboptimal because this requires removing cells from their native environment, mechanically or enzymatically disrupting tissue, and often exposing the cells to microfluidic manipulation. These processes can profoundly alter the metabolism of cells, especially microglia that are very responsive to their environment. Therefore, we measured metabolism in the M1 region of the murine cortex of fresh frozen brains taken from 16-week and 20-month male and female mice using FLIM to preserve the *in situ* metabolic profile of the brain as much as possible (**Figure 1A**) (Erkkilä et al., 2020; Han et al., 2023; Lukina et al., 2021; Ranjit et al., 2017; Sagar et al., 2020). Using the frequency domain FLIM approach we measured the fluorescence lifetime for nicotinamide adenine dinucleotide (NADH) and flavin adenine dinucleotide (FAD) simultaneously. The lifetimes of NADH and FAD are distinct depending on their function and microenvironment: lifetime of “free” fluorophore involved in processes in the cytoplasm is different from the lifetime of fluorophore “bound” to enzymes in the mitochondria and involved in oxidative phosphorylation (OXPHOS). Representative FLIM maps for free NADH and FAD are shown in **Supplemental Figure 1**. By determining the proportion of “free” and “bound” NADH, we were able to calculate the glycolytic index, with a higher ratio of free:bound indicating more glycolysis (**Figure 1B-C**). Analysis of FAD lifetime also allows us to determine the proportion of FAD that was free, versus the proportion that was “bound” to metabolic enzymes. By dividing the fraction of “bound” NADH by the fraction of “bound’ FAD, we were able to calculate an optical correlation of the cellular redox ratio: the fluorescence lifetime imaging redox ratio (FLIRR) (**Figure 1B-D**). This provides a measure of mitochondrial OXPHOS within a cell (Cao et al., 2019).

**Figure 1.**
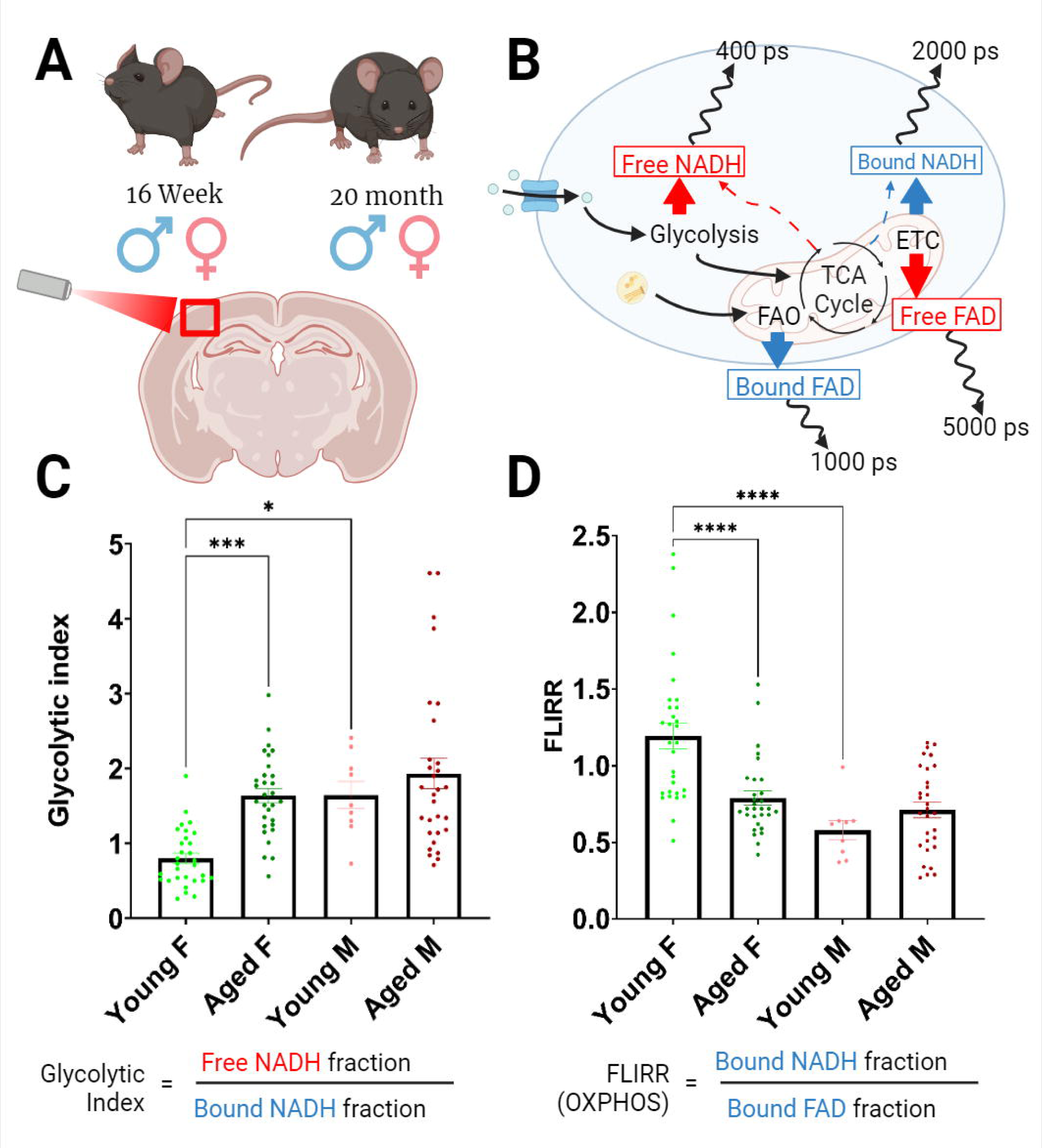
Fluorescence Lifetime Imaging Microscopy (FLIM) of young and old female brains. **A.** Schematic representation of model and sample preparation. The M1 region of the cortex was imaged using FLIM for all brain sections. **B.** Schematic representation of FLIM. **C.** Glycolytic indices of young (16-week) and old (20-month), female and male brain cortices showing increased glycolytic index with aging in females, with equation for calculations of glycolytic index. **D.** Fluorescence Lifetime Imaging Redox Ratio (FLIRR) of young and old, female and male brain cortices indicating less oxidative metabolism in the cortex of females with aging but more in young females than male counterparts with equation for FLIRR. No significant effects of aging were observed in males. (* p < 0.05; *** p < 0.001; **** p < 0.0001) Created with BioRender.

Interestingly, we found that the glycolytic index was higher in the young males than the young females (*p<0.05; **Figure 1C**). However, with aging the glycolytic index significantly increased in the females, such that the aged female brains exhibited a glycolytic index comparable to the young and aged male brains (***p<0.001; **Figure 1C**). Importantly, no significant changes were observed in the glycolytic index of males with aging. We also observed marked changes in the FLIRR of young males compared to young females, with females showing higher FLIRR, suggesting increased mitochondrial OXPHOS (****p<0.0001; **Figure 1D**). To gain further insight into mitochondrial metabolism with aging, we compared FLIRR in aged males and females (**Figure 1D**). We found that FLIRR was again unchanged in males with aging. However, in the female brains FLIRR was significantly reduced with aging, with values comparable to that of the male brains (****p<0.0001; **Figure 1D**). Therefore, these results suggest that with aging the metabolic profile of the female brain cortices appears to become more “male-like,” and highlight sex-specific changes in brain metabolism with aging, that may contribute to the metabolic reprogramming associated with AD.

### 2.2 Increased Expression of DAM and Metabolic Genes in Aged Female Microglia

Since our FLIM analysis revealed sex-differences in whole brain metabolism with aging, we next asked whether our findings could be explained by sex- and age-associated differences in microglial metabolism. We performed bulk RNAseq on CD11b+ microglia isolated from 16- week and 20-month female and male brains. Few genes were differentially expressed genes when comparing microglia isolated from young males and females (**Figure 2A-B**). However, significant differences were noted between sex-related genes such as *Xist*, *Ddx3y*, and *Eif2s3y* that are located on the X and Y chromosome (**Figure 2A**). Interestingly, in the 20-month mice, we observed significant differences in genes associated with AD pathology and disease-associated microglia (DAMs) (Olah et al., 2020), such as *Spp1*, *Clu*, and *ApoE* (de Luna et al., 2023; De Schepper et al., 2023; Kim et al., 2022) (**Figure 2B**). In addition, differential expression of X- and Y-linked genes like *Ddx3y*, *Uty*, *Kdm5d*, and *Eif2s3y* persisted with aging (**Figure 2B**).

**Figure 2.**
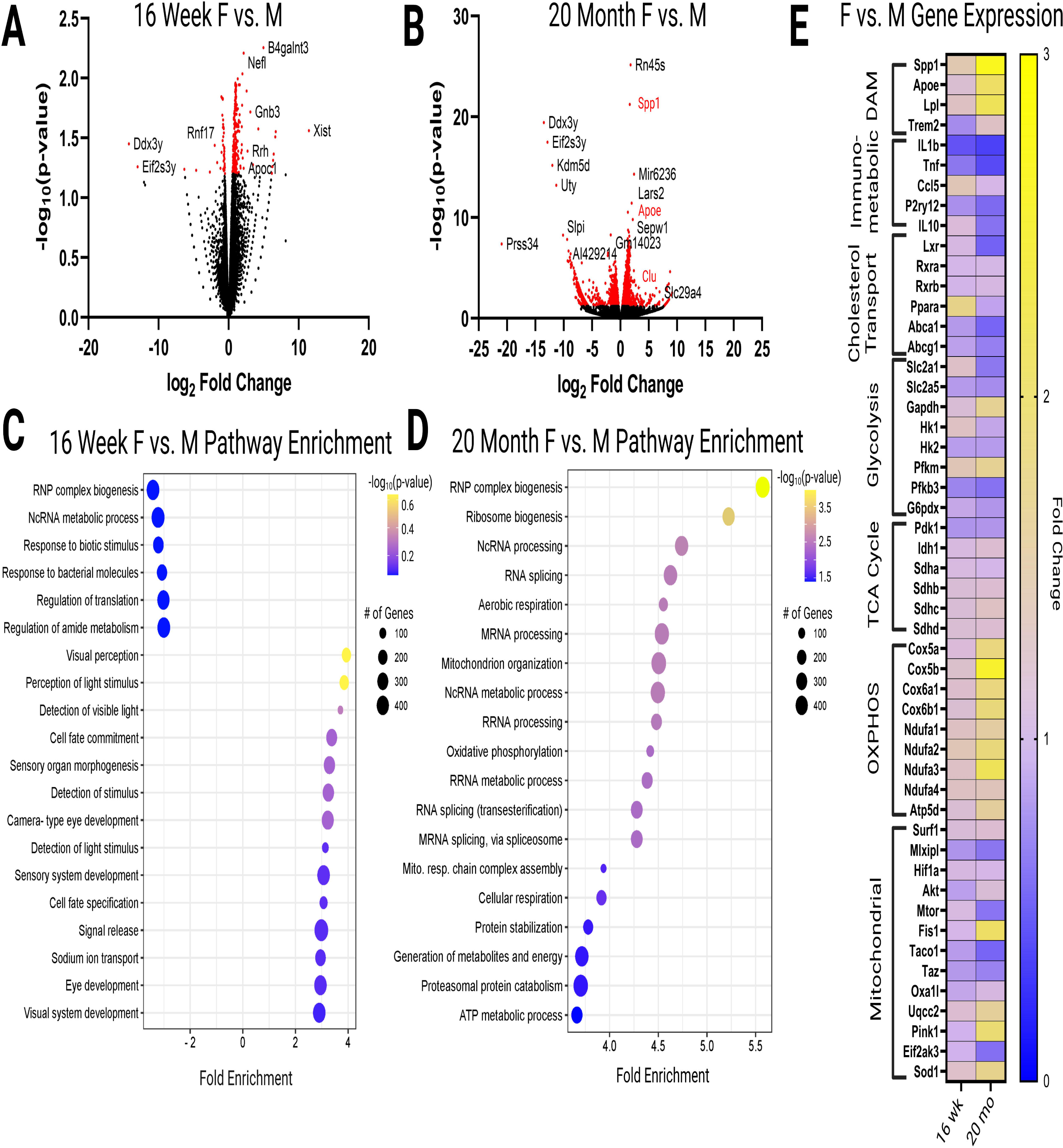
Differential gene expression of CD11b+ microglia isolated from 16-week and 20- month males and females reveals elevated metabolism and DAM signatures in aged females. **A**. Volcano plot comparing expression between 16-week males and females with significantly changed genes in red. **B**. Volcano plot comparing expression in 20-month males and females with significantly changed genes in red. **C.** Bubble enrichment plot showing differentially altered pathways in 16-week males and females. **D**. Bubble enrichment plot showing differentially altered pathways in 20-month males and females. **E.** Heatmap comparing specific disease-associated microglia (DAM), immunometabolic, cholesterol transport, glycolysis, TCA cycle, oxidative phosphorylation (OXPHOS), and mitochondrial genes in 16- week and 20-month females and males. Here, a decrease in enrichment is a value between 0-1. Created with BioRender.

We next performed a pathway enrichment analysis using differential gene expression data from 16-week and 20-month-old male and female microglia. Again, transcriptomic differences between 16-week male and female mice were minimal compared to the analysis of microglia from older mice. However, pathways such as “response to extracellular stimuli”, “signal release” and “sodium ion transport” appeared to be up-regulated in female microglia, whereas pathways such as “response to bacterial molecules” and “regulation of amide metabolism” were down-regulated in 16-week female microglia compared to male microglia (**Figure 2C**). Interestingly, comparative pathway analysis of microglial gene expression between 20-month female and male mice revealed significant age-related enrichment of metabolic pathways such as “aerobic respiration”, “oxidative phosphorylation”, “cellular respiration”, “protein catabolism”, and “ATP synthesis” in female microglia (**Figure 2D**).

To determine which specific genes may be driving changes in these metabolic pathways, we compared the expression of genes associated with metabolic processes critical to microglial function. First, we determined the expression of genes associated with lipid-processing and disease-associated microglia such as *Spp1, ApoE, LPL,* and *Trem2*. We found that expression of DAM genes was higher in females than males, and that this difference was more pronounced with aging (**Figure 2E**). In contrast, we observed downregulation of immunoregulatory genes (fold change <1), such as *TNF, IL-1*β, and *IL-10* in female microglia (**Figure 2E**). This change also appeared to become more pronounced with aging. Similarly, genes involved in cholesterol transport such as *Lxr, Ppara, Abca1,* and *Abcg1* were downregulated in female microglia, particularly with aging (**Figure 2E**). Regarding genes involved in transport of hexose sugars, we observed increased expression of the glucose transporter *Slc2a1* (GLUT1) in microglia isolated from young females compared to males (**Figure 2E**). Interestingly, with aging, expression appears to shift away from *Slc2a1* and towards *Slc2a5* (GLUT5), a fructose transporter associated with microglial activation and AD pathology (Burant et al., 1992; Tatibouët et al., 2000) (**Figure 2E**). Analysis of genes involved in glycolysis revealed increased expression of *Pfkm* and *Gapdh* in female microglia with aging (**Figure 2E**), which is consistent with recent studies highlighting upregulated glycolysis with aging in female microglia (Guillot-Sestier et al., 2021). Analysis of genes involved in mitochondrial function revealed increased expression of succinate dehydrogenase subunits (*sdha, sdhb, sdhc, sdhd*), Cytochrome C Oxidase (COX) subunits (*Cox5a, Cox5b, Cox6a1, Cox6b1*), NADH dehydrogenase subunits (*Ndufa1, Ndufa2, Ndufa3, Ndufa4*), and *Atp5d*, a subunit of adenosine triphosphate (ATP) synthase in female microglia with aging (**Figure 1E**). Although this suggests oxidative phosphorylation may be enhanced with aging in female microglia, analysis of other genes associated with mitochondrial dysfunction and oxidative stress (*Mlxip, Fis1, Pink1)* indicate that mitochondrial function may be dysregulated and/or uncoupled. Our findings are consistent with prior studies demonstrating exacerbated metabolic changes in female microglia with aging.

### 2.3 Estrogen alters DAM gene expression and lipid metabolism

Since sex-differences appear more pronounced with aging, when endogenous estrogen levels are dramatically reduced, we next asked how 17β-estradiol (E2) influences microglial metabolism. Here, we used murine BV-2 microglia. Although BV-2 microglia show altered production of cytokines compared to primary microglia (Angst et al., 2023; Henn et al., 2009), BV-2 cells are particularly useful for modelling AD, as they abundantly express DAM genes, such as *LPL* and *ApoE* (Loving et al., 2021; Oldham et al., 2022), unlike other microglial cell lines or primary microglia. Therefore, we asked whether E2 supplementation could alter the expression and function of DAM genes. Prior to conducting these experiments, we used RT-PCR to identify the sex of origin of BV-2 cells. We identified them as genetically female, and also confirmed that they expressed estrogen receptors (**Table 1**). Interestingly, we found that BV-2 cells predominantly express estrogen receptor 1 (ESR1 or ERα), consistent with our data from CD11b+ microglia isolated from male and female mice, and human microglia (Zhang et al., 2016) (**Table 1**). To ensure the absence of ESR2 expression was not due to the expression of tissue specific splice variants in BV-2 cells, we used a series of primers specific to other splice variants. We did not observe any amplification of transcripts associated with ESR2 splice variants in BV-2 microglia (**Table 1**).

**Table 1.** ESR1 and ESR2 Expression in microglia. BV-2 microglia are female, and only express ESR1. RNA sequencing studies using CD11b+ positive primary microglia show that murine microglia only express ESR1. Human microglia predominantly express ESR1. (Zhang et al., 2014; Zhang et al., 2016). (+ = expressed, ++ highly expressed, - = not expressed).

To determine the effect of E2 on BV-2 cell metabolism and phenotype, cells were supplemented with 100 nM E2 for 24 hours and then processed for downstream analyses, such as gene expression, LPL activity, and cholesterol efflux (Loving et al., 2021) (**Figure 3A**). Interestingly, following supplementation with 100 nM E2, the expression of tumor necrosis factor alpha (TNF- α) and key DAM genes *Trem2*, *ApoE*, and *LPL*, were reduced (**Figure 3B**). Since DAM genes were profoundly upregulated in aged female microglia (**Figure 2**), our data suggests E2 may be involved in negative regulation of a DAM gene cassette. We next measured the hydrolytic activity of LPL in BV-2 cells exposed to E2. Measuring extracellular and intracellular LPL activity is particularly important, since the enzyme is largely regulated at a post-translational level in many tissues (Doolittle et al., 1990; Shang and Rodrigues, 2021). In addition, comparing LPL activity within each cellular compartment indicates whether the enzyme has been exported to the cell surface to process lipids (Oldham et al., 2022). Typically, these metabolic assays are carried out in low serum conditions (1% fetal bovine serum [FBS]). We found that in low serum conditions, neither extracellular nor intracellular LPL activity was altered (**Figure 3C**). Considering gene expression analysis was performed in normal serum concentrations (10% FBS), and we observed a trend towards reduced LPL gene expression following E2 addition, we repeated these experiments in the presence of 10% FBS. We then found that intracellular LPL was increased in typical serum conditions following supplementation with E2 (**Figure 3C**). Our data suggest that E2 leads to intracellular retention of LPL during typical substrate availability but has less of an effect during low substrate availability. Since we have previously shown that reduced extracellular LPL leads to reduced cholesterol efflux (Loving et al., 2021), a process that is critical to microglial lipid droplet accumulation and lipid homeostasis, we next asked whether E2 supplementation could alter cholesterol efflux in BV-2 cells. We found that E2 supplementation reduced cholesterol efflux in BV-2 cells (**Figure 3D**). Moreover, we have also shown that the ESR1 specific agonist PPT (4,4’,4’’-(4-Propyl-[1*H*]-pyrazole-1,3,5-triyl) *tris*phenol) recapitulates the effects of E2 on cholesterol efflux, supporting the notion that E2- mediated changes in metabolism work predominantly through ESR1. Further, supplementation with the ESR2 specific agonist diarylpropionitrile (DPN), did not influence cholesterol efflux (**Figure 3D**). In further support, supplementation with PPT showed a trend towards E2-like effects on the expression of DAM genes in microglia (**Supplemental Figure 3**). Overall, these findings suggest that estrogen may down-regulate lipid processing in microglia, hence, lipid processing may be elevated in microglia of the aging brain where estrogen levels are reduced.

**Figure 3.**
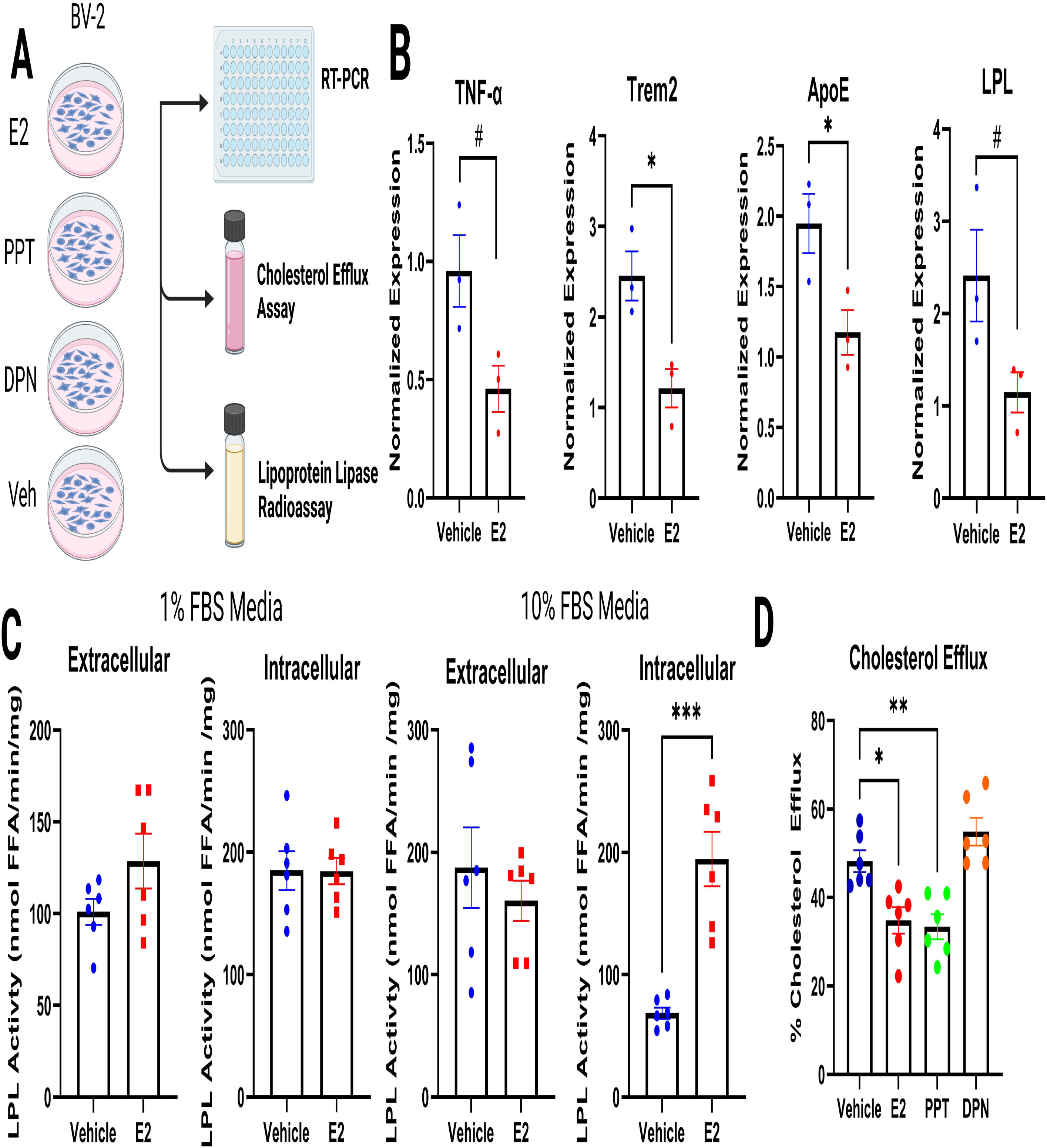
Estrogen acts via ESR1 to alter the expression of DAM genes and lipid metabolism. **A.** Schematic of experiments carried out in BV-2 microglia using ESR1 and ESR2-specific agonists 4,4’,4’’-(4-Propyl-[1*H*]-pyrazole-1,3,5-triyl) *tris*phenol (PPT) and diarylpropionitrile (DPN) respectively. **B.** qPCR gene expression data of *TNF-*α*, IL-1*β*, LPL, Trem2*, and *ApoE* in BV-2 microglia-like cells after 24 hours of drug exposure (estradiol (E2), or vehicle). Individual data points represent biological replicates. **C.** Lipoprotein lipase (LPL) activity in BV-2 microglia-like cells grown in either 1% FBS or 10% FBS exposed to E2 for 24 hours. Data points reflect biological and technical replicates. **D**. Cholesterol efflux from BV-2 cells exposed to E2, PPT or DPN 24 hours. Data points reflect biological and technical replicates. (# p < 0.1, * p < 0.05, ** p < 0.01, *** p < 0.001) Created with BioRender.

Since we observed increased LPL activity in E2-treated BV-2 cells (**Figure 3**), we asked if there were sex differences in the hydrolytic activity of LPL in whole brains. Here, brains were harvested from 9-month-old 5XFAD, a mouse model of AD that expresses 5 familial AD - mutations (Oakley et al., 2006), and age matched wild type (WT) mice. Intracellular and extracellular angiopoietin-like 4 (Angptl4)-sensitive LPL-specific lipase activity was quantified in whole brain tissue (Oldham et al., 2022). Notably, in the aged-mammalian brain LPL is predominantly expressed in microglia (Zhang et al., 2014). Hence, although there may be minimal contribution from other cell types, quantifiable LPL activity is largely from microglia. Interestingly, we found that extracellular LPL activity was significantly higher in WT female brains compared to males (**Figure 4A**). Notably, these sex-differences were not as pronounced in 5XFAD mice (**Figure 4A**). We also observed higher intracellular LPL activity in the brains of female WT mice compared to males (**Figure 4A**). While intracellular LPL activity was also higher in the brains of female 5XFAD mice, this difference was less pronounced than in the WT mice (**Figure 4A**). Taken together, this data shows for the first time to our knowledge that LPL activity is higher in aged female brains compared to males, and intriguingly may be linked to natural aging rather than accelerated disease status.

**Figure 4.**
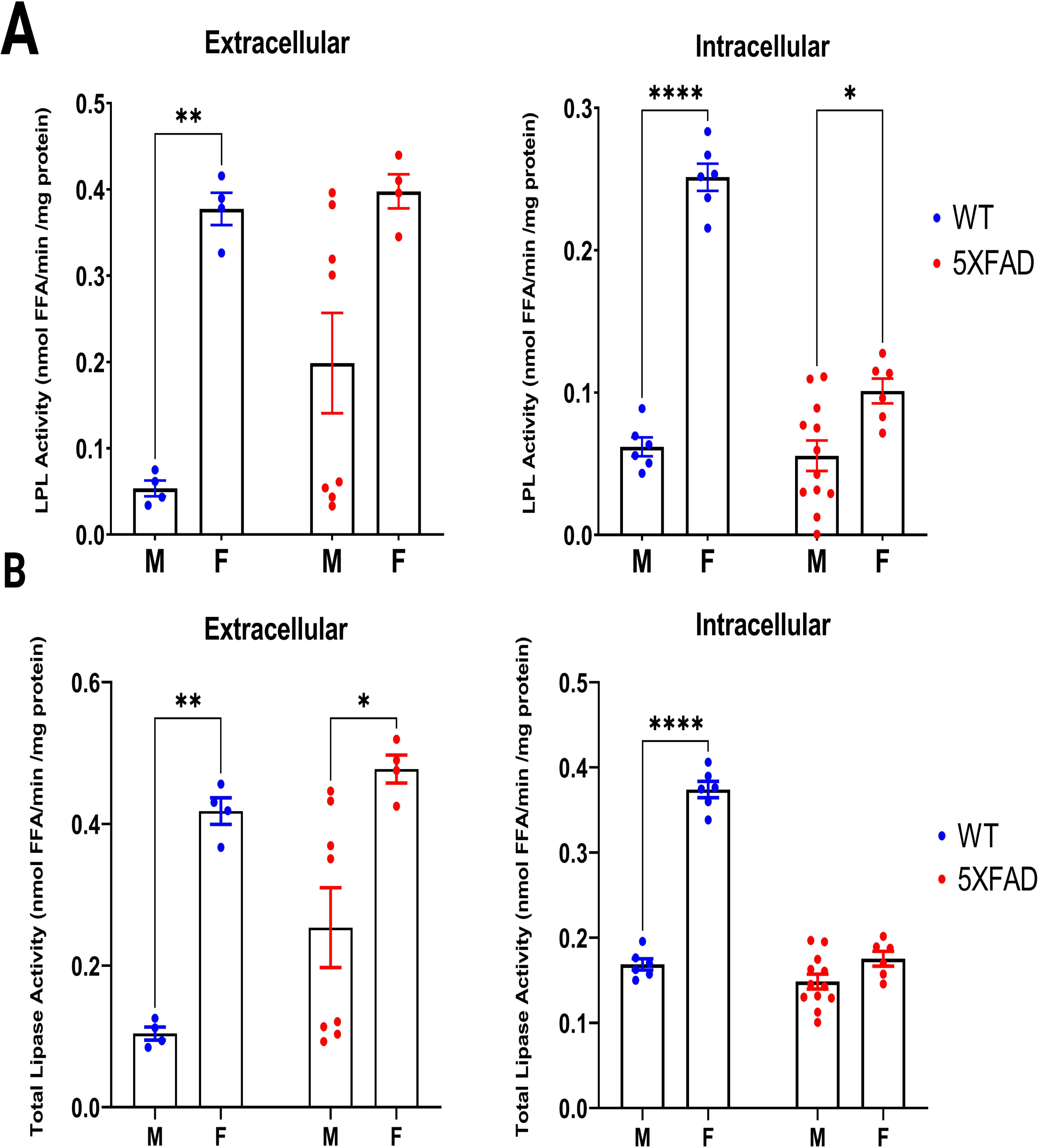
Lipoprotein Lipase (LPL) and Total Lipase Activity in aged Wild Type and 5XFAD Mice brains. **A.** Extracellular and intracellular LPL activity in whole brain tissue of 9- month-old wild type (WT) and 5XFAD male and female mice. (* p < 0.05; ** p < 0.01; **** p < 0.0001). Created with BioRender.

### 2.4 Estrogen may reduce Glycolysis and Increase Oxidative Metabolism in BV-2 microglia

Since E2 supplementation downregulated DAM gene expression and lipid metabolism in BV-2 microglia, we next asked whether E2 could also alter glycolysis and OXPHOS. We used FLIM to quantify the glycolytic index and FLIRR of BV-2 microglia exposed to 100 nM E2 for 24 hours. Since E2 supplementation may have substrate-dependent effects (**Figure 3**), we performed these experiments in both 1% and 10% FBS conditions. We found that in the 1% FBS condition, E2 did not significantly alter glycolytic index compared to vehicle (**Figure 5A**). Surprisingly, both vehicle and E2 (hence the presence of 0.001% ethanol) were sufficient to increase the glycolytic index of BV-2 microglia (**Figure 5A**). While this finding is somewhat tangential to the scope of this manuscript, it is important to consider the impact of ethanol, even as a vehicle, on microglial metabolism and function. In the 10% FBS condition, E2 supplementation, but not vehicle, decreased the glycolytic index of BV-2 microglia (**Figure 5A**). In contrast, E2 supplementation increased FLIRR in the 1% FBS condition (**Figure 5B**). Although a similar pattern was observed in the 10% FBS condition, the increase in FLIRR following E2 supplementation was not significant (**Figure 5B**). This provides further evidence for E2 treatment having differential effects on metabolism depending on substrate availability and nutrient status. To further investigate this, we conducted metabolomics on cells exposed to the same conditions. In the 1% FBS condition, glycolytic intermediate’s such as fructose-1,6- bisphosphate (F1,6BP), glyceraldehyde 3-phosphate (G3P), and 2- and 3-phosphoglycerate were significantly reduced following E2 supplementation (**Figure 5C**). We also observed increased glucose and pyruvate, the start and end products of glycolysis. This agrees with our FLIM data and indicates high glycolytic flux, during which we would expect low levels of glycolytic intermediates as they are quickly converted to the next glycolytic product, accompanied by high levels of start and end products. In the 10% FBS condition, we saw increased phosphoenolpyruvate (PEP) and 2- and 3-phosphoglycerate suggesting that glycolytic flux is reduced following E2 supplementation, consistent with the FLIM data (**Figure 5C**). Overall, these data suggest that E2 may downregulate glycolysis, but not in low FBS conditions, where lipids are limited, and the cells are more dependent on glycolysis to meet energy demands.

**Figure 5.**
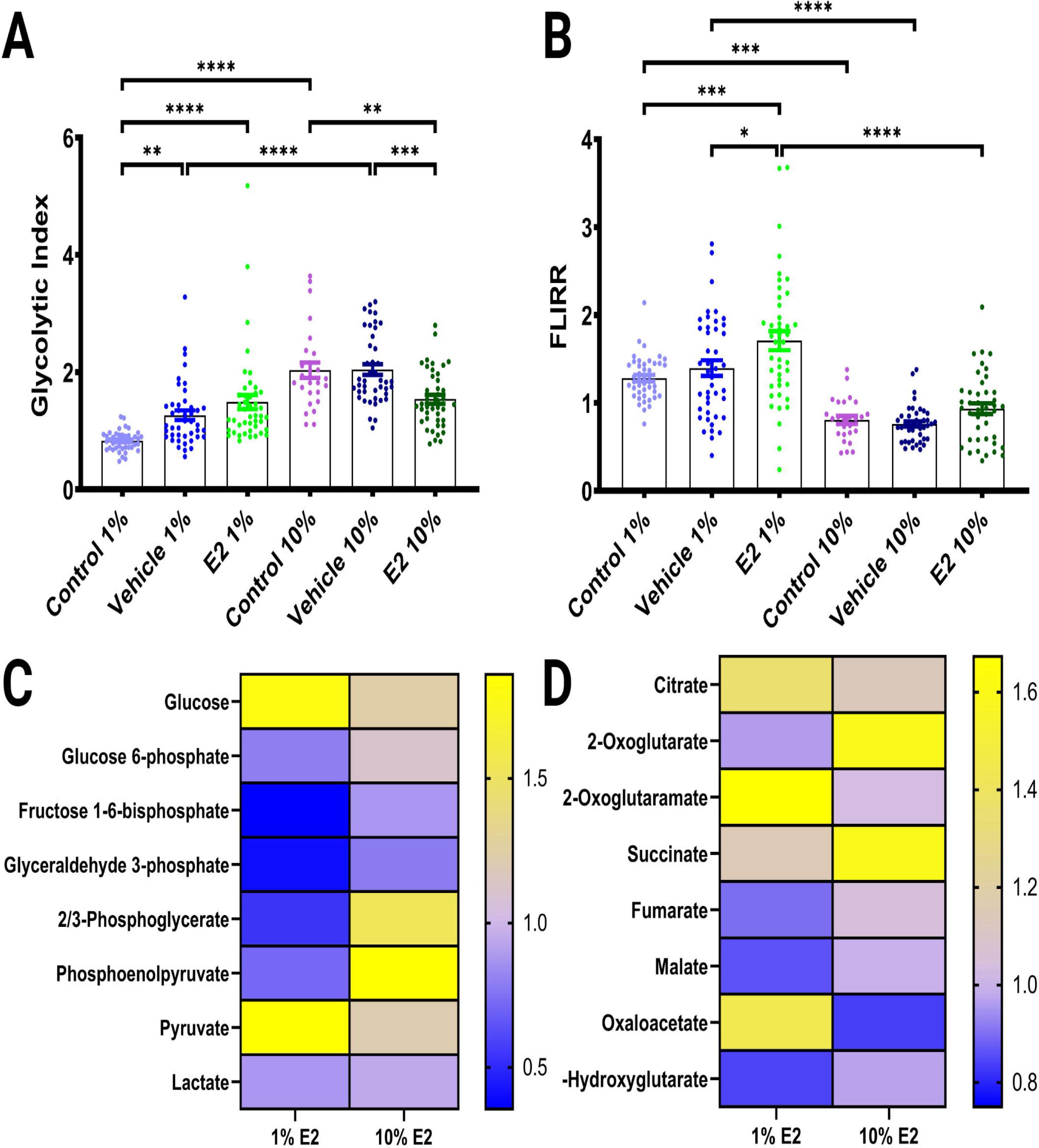
Metabolic effects of Estrogen on BV-2 microglia. **A.** Glycolytic index of BV-2 microglia in 1% or 10% FBS containing media treated with equal volumes of media (control), ethanol (vehicle), or E2 dissolved in ethanol (E2). **B.** FLIRR of BV-2 microglia in 1% or 10% containing media treated with equal volumes of media (control), ethanol (vehicle), or E2 dissolved in ethanol (E2). **C.** Heatmap of glycolytic intermediates from metabolomics data. **D.** Heatmap of TCA cycle intermediates from metabolomics data.

Looking next at the TCA cycle metabolites, we observed reduced fumarate, malate, and 2- hydroxyglutarate and increased citrate, 2-oxoglutaramate (2-OGM), succinate, and oxaloacetate (OAA) in E2 treated cells in the 1% FBS condition (**Figure 5D, Supplemental Figure 4**). A buildup of OAA suggests that citrate synthase may limit the TCA cycle, with a buildup of citrate suggesting that it is being shunted outside the mitochondria. This is also supported by the decrease in 2-oxoglutarate (**Figure 5D**). The increases in 2-oxoglutaramate and succinate suggest that the cells are breaking down glutamine and glutamate to further supply the TCA cycle with substrates (Hariharan et al., 2017). The increases in citrate, 2-oxoglutarate, and succinate in the 10% FBS condition could suggest that E2 supplementation enhances initiation of TCA cycle, but additional limiting factors (e.g., adenosine diphosphate [ADP]) may prevent completion (**Figure 5D**). Overall, however, the increase in TCA intermediates suggests that E2 supplementation increases the flux through mitochondrial metabolism, which is consistent with increased FLIRR in E2 treated cells.

### 2.5. Metabolic Effects of Estrogen in Human Microglia

Although BV-2 cells conveniently model microglia in an activated and DAM-like state, their cytokine and macrophage-like qualities limit their interpretation (Angst et al., 2023; Henn et al., 2009; McQuade et al., 2018). Recently, induced pluripotent stem cells (iPSCs), which can be differentiated into iPSC-induced microglia, have become more commonly used (Abud et al., 2017). However, these cells are expensive, and time-consuming to generate and maintain (Sheridan et al., 2022). Therefore, we adopted a microglia-like cellular model derived from peripheral blood monocytes (PBMCs) (McQuade et al., 2018; Sellgren et al., 2017). Here, PBMCs were differentiated into microglia-like cells using GM-CSF and IL-34 [134, 135]. In addition to differentiation, we added a maturation step supplementing cells with cluster of differentiation 200 (CD200) and chemokine ligand 31 (CXCL31) (McQuade et al., 2018). (See **Supplemental Figure 5**). After differentiation and maturation, the resulting monocyte-derived microglia-like (MDMi) cells abundantly expressed Triggering receptor expressed on myeloid cells 2 (Trem2), ApoE, and ionized calcium-binding adapter molecule 1 (Iba1) (See **Supplemental Figure 5**) and showed ramified Iba1+ staining (**Figure 6A**). Hence, the Blurton-Jones + maturation protocol was adopted for all MDMi experiments going forward (**Supplemental Figure 5**). Here, MDMi cells were cultured from a 55-year-old female and 54- year-old male donor. At the end of the differentiation and maturation process female cells were exposed to either vehicle or E2 for 48 hours. We found that *ApoE* expression was higher in male compared to female microglia and female microglia with E2 supplementation (**Figure 6B**). While this appears to contradict the notion that *ApoE* expression is increased in females with aging, we cannot control for prior metabolic and inflammatory signals that may also elevate *ApoE* expression in these cells in males. Interestingly, we found that *Trem2* expression was increased in middle-aged female MDMi cells compared to age-matched male counterparts, and that *Trem2* expression was reduced following E2 supplementation (**Figure 6B**). This is consistent with our findings in microglial cell lines and supports the notion that E2 supplementation can reduce DAM gene expression. We also observed a significant increase in interleukin-1beta (IL-1β) expression in female MDMi cells exposed to E2 when compared to males, with no differences observed between female MDMi cells exposed to vehicle vs. E2 (**Figure 6B**). Finally, *Slc2A5/GLUT5* expression was higher in females compared to males, although E2 supplementation appeared to reduce *Slc2A5* expression, this change was not significant (**Figure 6B**). Taken together, our findings suggest that there are immunometabolic sex-differences in microglia isolated from middle-aged females and males, and that these genes are also responsive to E2 supplementation.

**Figure 6.**
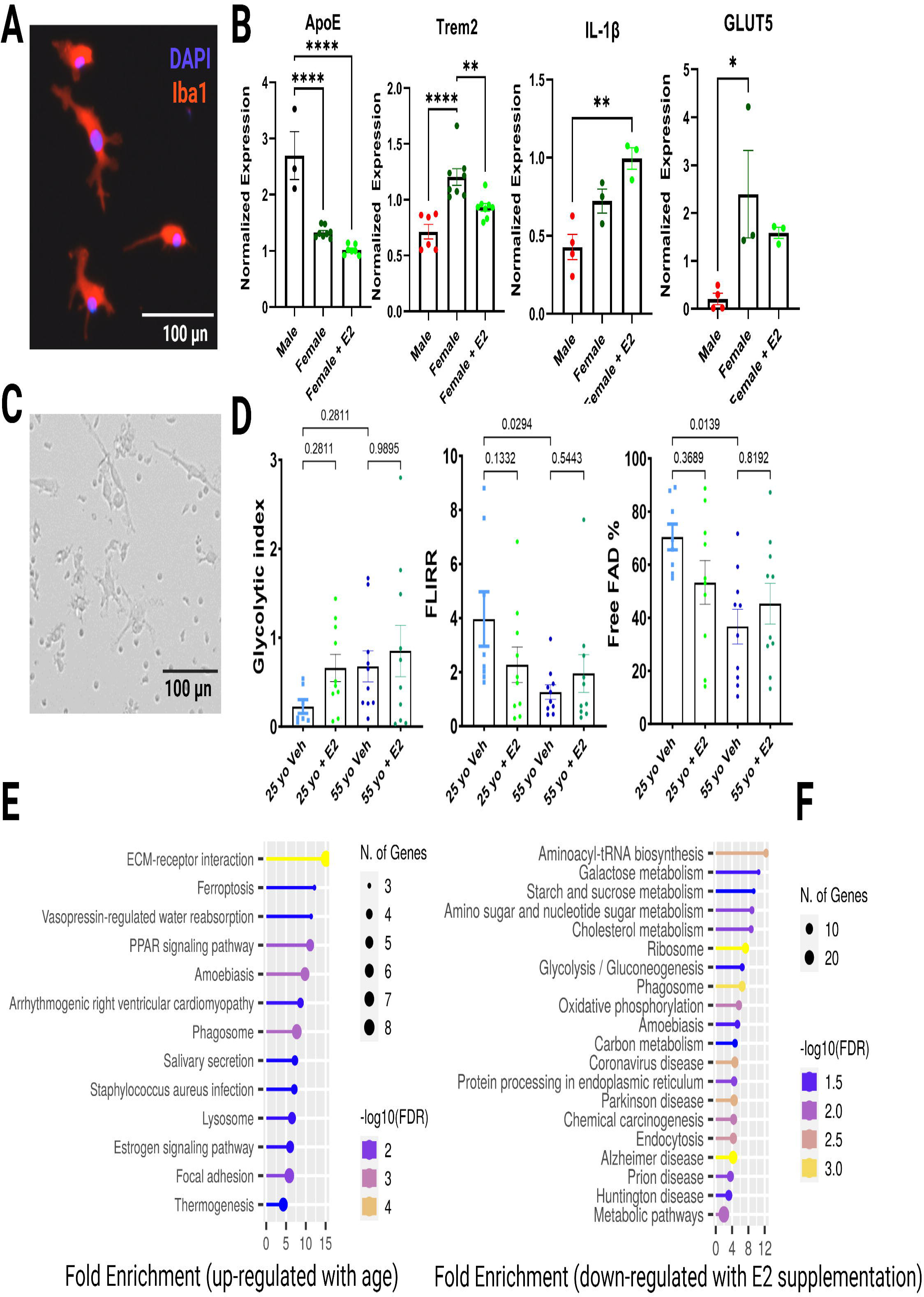
Metabolic effects of Estrogen on Human Microglia. **A.** Representative image of monocyte-derived microglia-like (MDMi) cells stained with DAPI (blue) and Iba1(red). **D**. Normalized expression data from MDMi cells from an aged male and female donor, with female MDMi cells treated with E2 for 48 hours. **C.** Representative brightfield image of MDMi before FLIM analysis. **D**. FLIM analysis of MDMi derived from 25 yo female or 55 yo female donors after treatment with vehicle (0.000001% Ethanol) or E2 for 48 hours. Individual data points reflect biological and technical replicates. (* p < 0.05; ** p < 0.01; *** p < 0.001; **** p < 0.0001).

Next, we used FLIM to determine age-associated metabolic changes in female MDMi, and to assess whether there was a metabolic response to E2 supplementation. After differentiation and maturation, microglia derived from young (25 year old) and middle-aged (55 year old) donors were supplemented with either vehicle or E2 for 48 hours. FLIM was used to measure glycolytic index (**Figure 6D**). Although there were no significant changes, there was a trend towards increased glycolysis with aging. In addition, the response to E2 supplement appears blunted with aging. FLIRR analysis revealed reduced oxidative metabolism in aged microglia, suggestive of mitochondrial dysfunction (**Figure 6D**). Consistent with this notion, free FAD was reduced in microglia derived from middle-aged compared to young donors (**Figure 6D**). Although E2 supplementation in the aged microglia appeared to move metabolic readouts to the direction of the young cells, this was not significant and further studies with additional donors are warranted to determine if E2 can restore or improve microglial metabolism and function in disease states.

To further determine changes in human microglial metabolism with age and E2 supplementation, we determined changes in media proteins from the MDMi cultures described above. KEGG pathway analysis revealed that extracellular matrix ‘(ECM)-receptor interactions’ and ‘PPAR signaling’ were up-regulated in young MDMi cultures compared to middle aged (**Figure 6E**). Notably, SPP1 (Osteopontin [OPN]) and LPL were among the main proteins driving the ECM and PPAR signaling pathways respectively (**Supplemental Figure 6A**), supporting the notion that DAM-associated factors increase with age in human and murine microglia. Interestingly, many of the protein pathways downregulated in middle-aged compared to young microglia were involved in metabolic processes such as the TCA cycle intermediates (**Supplemental Figure 6B**), consistent reduced mitochondrial function and metabolism with aging. In addition, analysis of media proteins revealed a marked reduction in many metabolic pathways upon supplementation with E2 (**Figure 6F**). For example, ‘cholesterol metabolism’, ‘Glycolysis’ and ‘Alzheimer’s disease’ pathways were downregulated in middle-aged MDMi cultures following E2 supplementation (**Figure 6F**). Analysis of the main proteins driving these pathway changes revealed down-regulation of APOC1 and LPL (cholesterol metabolism), HK1 (Glycolysis) and ADAM10, LRP1 and LPL (Alzheimer’s disease), which are among the top 20 downregulated proteins (**Supplemental Figure 6C and Supplemental Table 2**). Overall, these data suggest that while age may up-regulate microglial expression of proteins involved in lipid metabolism and AD pathogenesis, E2 supplementation may at least partially restore immunometabolic reprogramming in human microglia-like cells.

## 3. Discussion

Although AD is more prevalent in women, the mechanisms remain elusive. Here we have used a unique combination of metabolic imaging, transcriptomics, and functional measures of metabolism to show that metabolism is profoundly altered in the female brain and microglia, which may contribute to the increased risk of AD, and indeed other neurodegenerative diseases in women. Specifically, here for the first time to our knowledge we have used FLIM to show that with aging, the female brain becomes more glycolytic and less oxidative in females (**Figure 1**). Notably, no changes in glycolysis or oxidative metabolism were observed in males. This exacerbated change with aging in females likely contributes to altered brain function and increased disease risk in females. Our findings support recent studies that suggest that glycolysis is increased in the female brains with aging (Guillot-Sestier et al., 2021) and build on these findings by describing metabolic changes *in situ*. However, they are somewhat contradictory to observations reporting reduced glucose utilization with aging and disease. We reason that these inconsistencies are due to both regional differences, disease severity, and the contribution of glial metabolism. For example, Vaishnavi et al showed that aerobic glycolysis was elevated in the medial, lateral and prefrontal cortices of typical young adults at rest, but lower in the cerebellum and medial temporal lobes (Vaishnavi et al., 2010). Therefore, glycolysis appears to be higher in regions of the brain where Aβ accumulation is more prevalent. Hence, it is likely that increased glycolysis is a compensatory response to Aβ accumulation and mitochondrial dysfunction, which occur early in AD neuropathogenesis (Goyal et al., 2014; Goyal et al., 2020). In the later stages of AD, severe loss of neuronal function and brain atrophy eventually leads to reduced glucose utilization (Kim et al., 2005). Indeed, recent studies have suggested that altered glucose utilization in the brain may be an important predictor of AD outcomes (Goyal et al., 2020; Kim et al., 2005; Vaishnavi et al., 2010). Our findings suggest that age-associated changes in glucose utilization and mitochondrial function are more pronounced in females, which may highlight premature utilization of brain reserves, and a greater susceptibility to AD neuropathogenesis. However, numerous questions remain, and studies that use FLIM to determine brain metabolism in different regions and at different disease stages would help decipher the etiology of the metabolic changes associated with AD.

Since our FLIM analysis revealed changes in whole brain metabolism, we reasoned that changes in glial metabolism may underly the increased glycolysis with aging observed in females. This is supported by the notion that microglia increase glucose utilization when activated and during the development of neurodegenerative diseases such as AD (Guillot-Sestier et al., 2021). Surprisingly, our transcriptomic analysis of microglia isolated from young and old male and female brains revealed an increase in genes involved in glycolysis *and* oxidative phosphorylation in aged female microglia compared to males. Although this suggests oxidative metabolism may be enhanced with aging in female microglia, the expression of genes that support mitochondrial function were reduced, whereas genes associated with oxidative stress were elevated indicating mitochondrial function may be dysregulated and/or uncoupled (Manczak et al., 2004; Wang et al., 2014). For example, *Mlxipl*, a transcriptional activator involved in shunting glycolysis end products into the tricarboxylic acid (TCA) cycle or energy storage (Kooner et al., 2008), is reduced in 20 month female microglia. In contrast, *Fis1*, a gene involved in mitochondrial fission and a marker of AD (Wang et al., 2012), and, *Pink1*, a gene that is increased in Parkinson’s disease and oxidative stress (Cantuti-Castelvetri et al., 2007; Mei et al., 2009), both show upregulated expression in females with aging. It is also plausible that increased mitochondrial gene expression is a functional compensation relating to aging associated mitochondrial dysfunction and oxidative stress, repeatedly observed in the brain of AD patients (Manczak et al., 2004; Wang et al., 2014). Overall, our data suggest that metabolic reprograming in microglia is accelerated in female brains with aging, which may translate to increased susceptibility to AD.

One of the most robust changes observed in our transcriptomic analysis is the increase in DAM gene expression in female microglia with aging. DAM genes include *LPL, ApoE*, *SPP1*, and *Trem2* which have been associated with AD (Butovsky and Weiner, 2018), and hence include genes that are involved in brain lipid and lipoprotein processing. Considerable debate remains whether the elevation of these genes is a maladaptive functional compensation, or the appropriate response needed to maintain homeostasis in the aged and diseased brain. Nonetheless, the age-associated elevation in the DAM gene cassette is clearly exacerbated in microglia isolated from aged female brains (**Figure 2**). Since aging is associated with a rapid decline in circulating estrogen levels, our data suggests that estrogen may inhibit DAM gene expression. Although there are caveats in their utility, we used BV-2 microglia, which abundantly express DAM genes, to address this question. Interestingly, we found that E2 supplementation reduced the expression of *LPL, ApoE*, and *Trem2*; increased intracellular retention of LPL; and reduced cholesterol efflux (**Figure 3**). Importantly, our analysis of MDMi showed that DAM genes may also be elevated in middle-aged and post-menopausal females, and expression of these genes can be at least in part rescued by E2 supplementation (**Figure 6**). While studies in additional donors are needed, our findings in microglia isolated from aged mouse brains, microglia-like cells, and human MDMi support the notion that DAM genes are elevated with aging and may be suppressed with estrogen treatment.

While many studies have shown that DAM gene expression is elevated with aging (Coales et al., 2022; Riedel et al., 2016; Sala Frigerio et al., 2019; Sierksma et al., 2020), whether aging also elevates the function of the encoded proteins has not yet been determined. Here, for the first time we provide supportive evidence that active LPL is increased in aged female brains compared to males (**Figure 4**). The significance of intracellular LPL activity may need to be interpreted with caution, since intracellular LPL may not have access to endogenous LPL activators and may simply be stored in an active form in intracellular pools. However, heparin releasable extracellular LPL indicates active enzyme at the cell surface, consistent with our current model of LPL being tethered to the microglial membrane by heparin sulfate proteoglycans. Although we only performed these analyses with brains from 9-month-old mice, and analyses in younger and older mice are warranted, since estrogen levels rapidly decline in C57/BL6 mice by 8 months of age (Habermehl et al., 2022), our data suggest that estrogen may inhibit LPL activity in females, in addition to gene expression. Hence, LPL activity may be elevated in the context of declining E2 concentrations. This is consistent with previous studies showing that estrogen can negatively regulate LPL, albeit in a tissue specific manner (Hamosh and Hamosh, 1975; Homma et al., 2000; Iverius and Brunzell, 1988; Kim and Kalkhoff, 1975; Peinado-Onsurbe et al., 2008) Additionally, studies in ovariectomized mice have shown that E2 supplementation can reduce LPL activity and gene expression in adipose (Hamosh and Hamosh, 1975; Iverius and Brunzell, 1988; Kim and Kalkhoff, 1975; Peinado-Onsurbe et al., 2008). Moreover, this inhibition has been shown to be mediated by ligand-dependent estrogen receptor activity at a specific AP-1-like TGAATTC sequence located upstream from the LPL promoter (Homma et al., 2000) (Pedersen et al., 1991; Pedersen et al., 1992). Our data extend these findings into the murine brain, microglia-like cell lines and microglia derived from human cells, and suggest that the expression of *LPL, ApoE,* and *Trem2* are suppressed by E2 in an estrogen receptor-dependent manner. Notably, LPL activity is also higher in the brains of 9-month 5XFAD mice compared to control, but the sex-differences are less pronounced (**Figure 4**). Since LPL-expressing microglia appear to respond to Aβ accumulation in an attempt to wall-off Aβ plaques to prevent neurodegeneration, it is likely that LPL expression is higher in 9-month 5XFAD males (with abundant Aβ accumulation) compared to WT males, reducing the sex-differences in microglial LPL expression and activity. Notably, analysis of the secreted proteome of young versus old human microglia-like cells also revealed increased DAM associated factors such as LPL and SPP1, the expression of which was downregulated following E2 supplementation. Overall, it remains to be determined whether E2-mediated effects on microglial lipid processing precede or mitigate AD risk. Nonetheless, our findings highlight the potential of E2 to modify DAM-associated factors associated with microglial dysfunction and AD onset.

Our data suggest a possible role for hormone replacement therapy in menopausal women to restore the microglial metabolic reprogramming associated with AD neuropathogenesis. Although there have been several trials addressing the risks and benefits of HRT, they have yielded mixed results (Saleh et al., 2023; Wu et al., 2020; Zhang et al., 2021). Moreover, these treatments do not come without side effects, and it is often difficult to tease apart whether the benefits outweigh the risks. For example, estrogen increases thromboembolic risk, which can make it more difficult to discern vascular dementia from Alzheimer’s dementia (Vinogradova et al., 2019). Systematic reviews evaluating data from these studies highlight the need for additional studies to determine whether HRT’s benefits outweigh the risks (Wu et al., 2020; Zhang et al., 2021). However, it is thought that if HRT is started early enough within the critical window it can be beneficial for AD (Wu et al., 2020). In addition, a recent study showed that HRT may have a greater benefit for women that carry the AD risk gene ApoE4 (Saleh et al., 2023). Since ApoE4 is primarily a lipid transport protein, it highlights the role of estrogen in regulating lipid metabolism in microglia and cells for further studies that address whether estrogen can specifically address the metabolic reprogramming observed in ApoE4. Notably, although data supporting the role of follicle stimulating hormone (FSH) antagonism appears promising for the treatment of AD (Xiong et al., 2022), we did not focus on the role of FSH, since microglia do not express FSH receptors (Zhang et al., 2014; Zhang et al., 2016).

The estrogen receptor expression profile of microglia and the CNS has been debated over the last few decades and remains an area of active investigation (Gold et al., 2009; Qu et al., 2022; Saijo et al., 2011; Wu et al., 2013). The data we present here relies on both qPCR data using primers designed to span an exon-exon junction, as well as nuclear ESR agonists with very high specificity (Sepehr et al., 2012). Our data contradicts previous studies that have found that BV-2 microglia expressed ESR2 (Baker et al., 2004), but is consistent with our *in vivo* data and published studies that have reported ESR1 expression in mouse and human microglia (Zhang et al., 2016).Our data using ESR1 specific agonists (e.g., PPT), which recapitulate our findings regarding E2 mediated changes in gene expression and cholesterol efflux (**Supplemental Figure 3**), further support the notion that ESR1 is the predominant (but not only) estrogen receptor in microglia. Of note, we did not focus on the role of the G-protein coupled estrogen receptor (GPER), and it is possible that some of the effects we found are mediated through this surface receptor (Guan et al., 2017; Qu et al., 2022). Indeed, other research has suggested that the release of proinflammatory cytokines like interleukin-1β (IL-1β) and tumor necrosis factor α (TNF-α) are mediated by GPER (Guan et al., 2017). Overall, our data support the need for more research into the effects of selective estrogen receptor modulators like tamoxifen, raloxifene, and bazedoxifene and for the development of similar new drugs with cell-specific, and hence microglia-specific, effects.

While our study highlights exacerbated immunometabolic changes in female microglia with aging as a potential mechanism contributing to the increased susceptibility of AD in women, we acknowledge several caveats in our study design. While BV-2 microglia-like cells are convenient samples for studying metabolism, especially lipid metabolism, they are not a perfect representation of AD. While MDMi cells derived from PBMCs are of human origin rather than mouse, they too come with caveats. These cells come from specific donors, each with their own specific transcriptomic profile, epigenetic profile, differing disease susceptibility, and prior or current drug exposure to name a few confounding factors. This system is ideal for patient-specific models, but as a result lack generalizability. Additionally, several studies have demonstrated that PBMC cytokine production is affected my menopausal status, age, and nutritional state (Kim et al., 2012; Lee et al., 2017; Paik et al., 2012). Although they provide a useful model for investigating human microglia without requiring the time or resources needed to grow and reprogram iPSCs, follow-up studies are needed using additional donors at different ages and stages of disease, and different genotypes (e.g., ApoE4) to recapitulate these findings and to determine whether estrogen exposure or agonism can improve microglial function and AD outcomes in women.

## 4. Summary and Conclusion

Despite AD disproportionately affecting women, the mechanism for this is not fully understood. In AD, microglia undergo ‘metabolic reprogramming’, but sex-specific changes in microglial metabolism have been understudied. Here, we used metabolic imaging, transcriptomics, and metabolic assays to show that female brains exhibit accelerated metabolic dysfunction with aging. Moreover, transcriptomic analysis of microglia isolated from 16-week and 20-month-old mice revealed increased expression of DAM genes (e.g., *ApoE, Spp1, LPL*). E2 supplementation resulted in reduced expression of DAM genes, intracellular retention of LPL, and reduced cholesterol efflux. Analysis of MDMi cells revealed higher DAM and GLUT5 gene expression in middle-aged females compared to males, which may be responsive to E2 supplementation. Overall, our findings corroborate and extend previous studies and show that altered metabolism defines age-associated changes in female microglia that could increase AD risk and severity. Our work supports the need for further studies aimed at restoring microglial metabolism, potentially through hormone replacement therapy, to improve outcomes for women with or at risk of AD.

## 5. Materials and Methods

### 5.1 Animals Wild type and 5XFAD

All investigations using either WT (C57BL/6 [#000664-JAX]) or 5XFAD (#034840-JAX) animals in this study were carried out using protocols (Ref #0115 and #0815) that were approved by the University of Colorado Institutional Animal Care and Use Committee (IACUC), ensuring compliance with the recommendations in the Guide for the Care and Use of Laboratory Animals, Animal Welfare Act and PHS Policy. Mice were housed no more than 5 mice per cage and maintained at ∼20°C with a 12-hour light/dark cycle and given unrestricted access to standard laboratory diet (Diet 8640; Harlan Teklad, Madison, WI, USA) and water. At the end of each experiment mice were trans-cardially perfused with Hank’s Balanced Salt Solution (HBSS) (with calcium and magnesium), and brains were isolated and fresh-frozen in Tissue Tek using a liquid nitrogen isopentane bath. Notably, fresh frozen brain sections were generated not more than a few weeks prior to FLIM analysis.

### 5.2 BV-2 Microglia-like Cell Culture

BV-2 microglia-like cells (Timmerman et al., 2018) were cultured at 37 °C with 5% CO_2_ in a solution comprising Dulbecco’s modified Eagle’s medium (DMEM) with 10% heat-inactivated fetal bovine serum (FBS) and 1% penicillin/streptomycin (P/S). The cells were grown to ∼75– 90% confluency prior to drug exposure or functional assay for all experiments.

### 5.3 BV-2 Microglia-like Cell Drug Exposure

After cell culture had reached 75-90% confluency as described above, media was changed to include test conditions. Test conditions included 850 pM diarylpropionitril (DPN), 20 nM 4,4’,4’’-(4-Propyl-[1*H*]-pyrazole-1,3,5-triyl)*tris*phenol (PPT), 100 nM estradiol (E2) all in either 1% FBS or 10% FBS with 1% penicillin/streptomycin in DMEM with 4.5 g/L L-glucose. Drugs were dissolved in ethanol.

### 5.4 MDMi Cell Culture and Differentiation

MDMi cell culture and differentiation protocol was adapted from previously published methods (McQuade et al., 2018; Sellgren et al., 2017). PBMCs were isolated from human blood using the SepMate PBMC isolation tubes (StemCell) according to manufacturer’s instructions. PBMCs were then frozen in 90% FBS with 10% dimethyl sulfoxide (DMSO). Vials of frozen PBMCs were thawed in a 37 °C water bath for approximately 2 minutes prior to resuspension and washed in warmed media composed of RPMI-1640 (Gibco) supplemented with 10% heat-inactivated fetal bovine serum (Gibco), 1% P/S, 1% GlutaMAX (Gibco). Cells were then plated onto Geltrex (Gibco) coated 6-well plates at approximately 5 million cells and 2 mL of media per well of a 6-well plate or 1 million cells per well and 1 mL of media per well of a 24-well plate and cultured at 37 °C with 5% CO_2_. The following day, media was aspirated off and replaced with warmed RPMI-1640 media supplemented with 1% P/S, 1% GlutaMAX, 100 ng/mL IL-34, 25 ng/mL M-CSF, 50 ng/mL TGF-β (hereafter referred to as “Differentiation Media” from here on out). On every third day, half of the Differentiation Media was removed and replaced with the same quantity of new, warmed media containing cytokines at double their concentration in differentiation media. On day 10-13, all media was removed and replaced with fresh maturation media containing CD200 100 ng/mL and CXCL31 100 ng/mL. On day 13-15, cells were treated with either 10 nM E2 or vehicle (water). Timing for the transition from differentiation media to maturation media depended on appearance of the cells in culture (relative ramification, number of floating cells, amount of cell debris). For Iba1 staining, cells were fixed in 4% paraformaldehyde for 20 minutes, permeabilized with 0.1% Triton-X (ThermoFisher Scientific) and incubated with Iba1 mouse monoclonal primary antibody (Millipore MABN92) overnight at 4° C. Cells were imaged using a fluorescent microscope (NIKON).

### 5.5 Isolation of Microglia from Adult Mouse Brain

Adult mice were perfused with HBSS without calcium and magnesium. Half of each adult mouse brain was minced with a sterile razor blade and placed in papain solution (Worthington, Lakewood, NJ, USA) #LK003150, prepared according to manufacturer’s instructions) and shaken at 100 rpm for 30 min at 37 °C. Tissue was triturated three times with a 10 mL, 5 mL, and 1 mL (pulled glass) pipette, respectively, filtered with a 70 μm cell strainer, and resuspended in Earle’s Balanced Salt Solution with DNAse I. Dissociated cells were separated from myelin debris using the Worthington ovomucoid gradient and resuspended in PBS. CD11b microbeads (Miltenyi Biotech Inc, Auburn, CA, USA) were incubated with the cell suspension at 4 °C for 15 min. The cell suspension was added to a magnetic cell separation (MACs^®^) column (Miltenyi Biotech) and washed several times to remove non-CD11b^+^ cells. CD11b+ microglia were collected and immediately frozen in Trizol™ solution prior to RNA extraction.

### 5.6 RNA sequencing

Total RNA was isolated from isolated microglia using the Trizol-chloroform RNA extraction method. Equal amounts of RNA were pooled from 3 male and female brains at 16 weeks. 3 male and 3 female 20-month mouse samples were extracted and sequenced individually. A mean FPKM value was used for differential expression analysis. The Agilent Tape Station 4200 RK6 screen tape was used to assess the integrity of the Total RNA. 100 ng of Total RNA was used as input to construct mRNA libraries using the Nugen Universal Plus mRNA library prep kit catalog No. 0508. Library quality was assessed on a D1000 screen tape using the Agilent Tape Station 4200. Libraries were quantitated using the Qubit, diluted to 4 nM, and sequenced at a depth of 80 Million Paired End Reads 2 × 150 on the Illumina NovaSEQ6000 sequencer. bcl files were converted to FASTQ files using CASAVA 2.0 software. The RNA-Seq data was analyzed using Basepair software (https://www.basepairtech.com/) with a pipeline that included the following steps. Reads were aligned to the transcriptome derived from UCSC genome assembly (hg19) using STAR (Dobin et al., 2013) with default parameters. Read counts for each transcript was measured using featureCounts (Liao et al., 2014). Differentially expressed genes were determined using DESeq2 (Love et al., 2014) and a cut-off of 0.05 on adjusted p-value (corrected for multiple hypotheses testing) was used for creating lists and heatmaps, unless otherwise stated. GSEA was performed on normalized gene expression counts, using gene permutations for calculating p-value.

Graphs of RNAseq data were generated using GraphPad Prism 9th Edition. Pathway analysis was completed using iDEP.96 software (http://bioinformatics.sdstate.edu/idep/). Pathways were analyzed in iDEP.96 using the fold change and corrected p-value data derived from Basepair analysis. Pathway enrichment was determined using gene set enrichment analysis (GSEA) and cap analysis of gene expression (CAGE), employing both GO functional categorizations and KEGG metabolic pathways to characterize enrichment. Results were plotted as bubble enrichment plots using https://www.bioinformatics.com.cn/en, a free online platform for data analysis and visualization.

### 5.7 Quantitative Polymerase Chain Reaction

**(qPCR)** Total RNA was isolated from primary and BV-2 cells and MDMi cells using the Trizol-chloroform extraction method. Verso cDNA synthesis kit (Thermo Scientific, Waltham, MA, USA) was used to synthesize cDNA from RNA. Using housekeeping genes, 18S and UBC, to establish expression standards, gene expression levels relative to these standards were quantified by qPCR using SYBR Select Master Mix (Applied Biosystems, Foster City, CA, USA). The NIH sponsored NCBI primer-blast tool (https://www.ncbi.nlm.nih.gov/tools/primer-blast) was used to design primers (**Supplementary Table 1**) that crossed exon boundaries when possible. Using StepOnePlus instrument and software v2.3 (Applied Biosystems, Foster City, CA, USA), thermal cycling conditions for all qPCR studies were as follows: initial temperature of 50 °C for 2 min, 95 °C for 10 min, then 40 cycles of 95 °C for 15 s and 60 °C for 1 min.

### 5.8 Genotyping

Genotyping for the 5XFAD and mice was conducted following the Jax protocol for standard PCR genotyping(https://www.jax.org/Protocol?stockNumber=006554&protocolID=31769). Briefly, each sample consisted of 2 uL DNA, 6 uL water, 10 uL 2x GreenTaq Master mix, 1 uL forward primer and 1 uL reverse primer. This was run on a thermocycler before being loaded onto a 2% agarose gel along with a DNA ladder and run at 100 V for 1 hour. For mouse samples, tail snips were taken from mice and digested in a tail digestion solution consisting of 75 uL of a 50 mM 1M Tris HCl pH 8.0, 10 mM EDTA, 100 mM NaCl, 0.1% SDS solution, and 2 uL of Proteinase K (20 mg/mL). A similar protocol was conducted for sex genotyping of BV-2 cells and mice. Briefly, a master mix of 6uL water, 10 uL GoTaq Green master mix, and 1 uL of each primer at 10 μM was added to 2 uL of DNA. Samples were then run on a thermocycler with the following program: 30 cycles of 94 °C for 2 minutes, 94 °C for 20 seconds, 60 °C for 20 seconds, 72 °C for 30 seconds. After 30 cycles, 72 °C for 5 minutes. Samples were then run on a 1% agarose gel with a DNA ladder for reference.

### 5.9 Fluorescence Lifetime Imaging Microscopy

**(FLIM)** FLIM was performed to detect local metabolic changes in fresh frozen whole brain coronal sections (10 µm) and BV-2 microglia cells after 24-hour treatment with 100 nM E2 or vehicle (EtOH) using a Zeiss 780 laser-scanning confocal/multiphoton-excitation fluorescence microscope with a 34-Channel GaAsP QUASAR Detection Unit and non-descanned detectors for 2 photon fluorescence (Zeiss, Thornwood, NY, USA) equipped with an ISS A320 FastFLIM box and a titanium:sapphire multiphoton Mai Tai laser (Spectra-Physics, Milpitas, CA). The 2 photon excitation was blocked by a 2 photon emission filter. For the acquisition of FLIM images, fluorescence for nicotinamide adenine dinucleotide (NADH) and flavin adenine dinucleotide (FAD) was detected simultaneously by two photon-counting PMT detectors (H7422p-40; Hamamatsu, Japan). Images of the 15-20 different areas of cortex in the brains were obtained with VistaVision software (ISS, Champaign, IL, USA) in 256 × 256 format with a pixel dwell time of 6.3 µs/pixel and averaging over 30 frames. Calibration of the system was performed by measuring the known lifetime of the fluorophore fluorescein with a single exponential decay of 4.0 ns (Berezin and Achilefu, 2010). The phasor transformation and the data analysis were carried out using Global SimFCS software (Laboratory for Fluorescence Dynamics (LFD), University of California, Irvine) as described previously (Digman et al., 2008). The number of pixels covered with lifetimes for free and bound reduced forms of NADH and FAD were calculated in SimFCS (LFD), and the values were normalized to the total number of pixels detected as previously described (Dobrinskikh et al., 2019; Marwan et al., 2019). The glycolytic index was calculated for all experimental groups using the following equation, as defined previously (Dobrinskikh et al., 2019; Wallrabe et al., 2018):

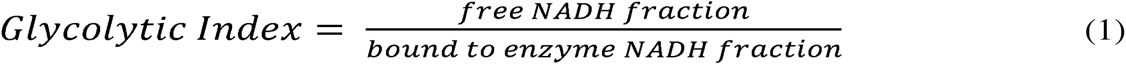

To assess OXPHOS and ergo mitochondrial activity, FLIM-based optical redox ratios (i.e. fluorescence lifetime redox ration (FLIRR)) was calculated as follows, as described previously (Dobrinskikh et al., 2019; Marwan et al., 2019):

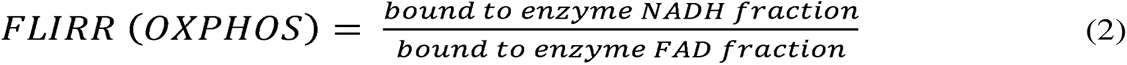

### 5.10 Cholesterol Efflux Assay

BV-2 microglia were seeded in DMEM (10% FBS), at a density of 75 K per well of a 12 well plate. The cholesterol efflux assay was adapted from (Low et al., 2012). When the cells had adhered to the plate (around 1 h), [^3^H]Cholesterol (Perkin Elmer) containing media was added to a final concentration of 0.5 μCi per well for 24 h. [^3^H]Cholesterol-containing media was removed and the cells were washed three times in sterile PBS. Serum-free DMEM was added to the cells, with T0901317 (Gibco) at 0–4 μMol/L for 20 h. Media was removed and 0.1 mL was added to 5 mL of Scintillation fluid and considered as “media counts”. The cells were washed three times with PBS and 0.5 mL of double-distilled water was added to the plates, which were then placed in a −80 °C freezer for one hour to help cells detach. After thawing, 0.1 mL of cell solution was added to 5 mL of scintillation fluid and the reads considered as “cell counts”. Cholesterol efflux was determined using the following equation:

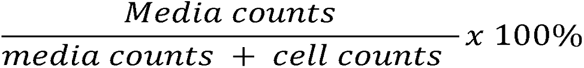

and expressed as a percentage (%) of total.

### 5.11 LPL Assay - Cell culture

LPL specific radiometric activity assays were completed as previously described (Oldham et al., 2022). In brief, cells were cultured as described above in 6 cm^2^ plates until at least 80% confluent. Cells were washed with HBSS with calcium and magnesium twice to remove any proteins from the media. Calcium and magnesium are crucial to prevent release of LPL from the cell surface. A Krebs Ringer Phosphate (KRP) buffer containing 80 ng/mL heparin sodium was added to the cells at 400 µL and incubated for 5 minutes at room temperature. The heparin binds to the surface LPL and releases it from the surface resulting in the KRP-heparin solution (KH) containing extracellular LPL. The KH was carefully transferred to an Eppendorf tube and kept on ice until use. The cells were washed and a second buffer of 80 ng/mL heparin in a buffer of 50% KRP and 50% Mammalian Protein Extraction Reagent (M-PER) was added to the cells. The cells were then scraped, and the lysis-heparin buffer (LH) was transferred to an Eppendorf tube with a single bead within it. The cell lysate was then shaken in a bead homogenizer for 2 minutes at 30 Hz to ensure complete lysis. The lysed cells were used to quantify intracellular LPL. LPL activity in each sample was determined by incubating each sample with a radiolabeled-triolein and ApoC-II containing substrate as previously described (Oldham et al., 2022). To ensure LPL specific activity, the LPL inhibiter recombinant human angiopoietin-like 4 N-Terminal fragment (Angptl4) was added to an extra replicate. The remaining activity (non-LPL) was subtracted from other technical replicates to estimate LPL specific activity.

### 5.12 LPL Assay - Tissue

LPL specific radiometric activity assay for tissue follows largely the same steps as for cell culture with a few differences. Briefly, frozen brains were thawed, then weighed. Brains were then minced with a razor blade and suspended in 1 mL of HBSS with calcium and magnesium before being passed through a 40 μm filter. The solution was then centrifuged at 500 g for 10 minutes, the supernatant discarded, and the pellet resuspended in 450 μL KH and let to incubate for 10 minutes at room temperature. The solution was centrifuged at 500 g for 5 minutes and the supernatant removed and kept on ice. 600 μL of LH and 1 zirconium bead were added to each brain, and the solution homogenized with a bead beater at 30 hz for 2 minutes. The solution was then centrifuged at 16,000 g for 20 minutes, and the supernatant removed and kept on ice. LPL activity in each sample was determined by incubating each sample with a radiolabeled-triolein and ApoC-II containing substrate as previously described (Oldham et al., 2022). To ensure LPL specific activity, the LPL inhibitor recombinant human angiopoietin-like 4 N-Terminal fragment (Angptl4) was added to an extra replicate. The remaining activity (non-LPL) was subtracted from other technical replicates to estimate LPL specific activity.

### 5.13 Metabolomics

BV-2 microglia-like cells described above were grown to 70% confluence prior to treatment with 100 nM E2 or vehicle (EtOH) for 24 hours. Each treatment condition was analyzed in triplicate. Frozen cell pellets were extracted at 2 × 10^6^ cells/mL in ice-cold lysis/extraction buffer (methanol:acetonitrile:water 5:3:2 v/v/v). Ultra HPLC (UHPLC)-MS-based high throughput metabolomics was performed at the University of Colorado School of Medicine Metabolomics Core. Metabolites were separated using a 9 min C18-based gradient method as previously described (McCurdy et al., 2016), using a Thermo Vanquish UHPLC coupled to a Thermo Q Exactive mass spectrometer. In brief, extracts (10 μL) were resolved in a Kinetex C18 column using a 3 min isocratic gradient at 250 μL/min (mobile phase: 5% acetonitrile, 95% 18 MOhm H_2_O, 0.1% formic acid) or a 9 min gradient (5% B for the first 2 min, 5%–95% B over 1 min, hold at 95% for 2 min, 95%–5% B over 1 min, re-equilibrate for 3 min). Quality control was performed via the assessment of a technical mix injected after every 10 samples as well as by comparison of internal standards. Metabolite assignments were performed with the software MAVEN (Melamud et al., 2010) upon conversion of .raw files into the mzXML format using MassMatrix). Assignments were further confirmed by chemical formula determination from isotopic patterns, accurate intact mass, and retention time comparison against an in-house standard library (Sigma-Aldrich, IROA Technologies, Sea Girt, NJ, USA). Pathway analysis and analysis of metabolite set enrichment was performed using MetaboAnalyst (Xia and Wishart, 2010a; Xia and Wishart, 2010b).

### 5.14 Proteomics

Media from 25 and 55 yo MDMi was pooled and centrifuged to remove non-adherent or dead cells. Samples were concentrated with a speedvac to ∼200ul and protein concentration was determined on a Nanodrod using the Absorbance 280 program. Plasma samples were digested in the S-Trap 96-well plate (Protifi, Huntington, NY) following the manufacturer’s procedure. Briefly, around 50 μg of plasma proteins were first mixed with 5% SDS. Samples were reduced with 5 mM tris (2-carboxyethyl) phosphine (TCEP) at 55°C for 30 min, cooled to room temperature, and then alkylated with 20 mM 2-chloroacetamide in the dark for 30 min. Next, a final concentration of 1.2% phosphoric acid and then six volumes of binding buffer (90% methanol; 100 mM triethylammonium bicarbonate, TEAB; pH 7.1) were added to each sample. After gentle mixing, the protein solution was loaded to an S-Trap 96-well plate, spun at 1500LJ*g* for 2 min, and the flow-through was collected and reloaded onto the 96-well plate. This step was repeated three times, and then the 96-well plate was washed with 200 μL of binding buffer three times. Finally, 1 μg of sequencing-grade trypsin (Promega) and 125 μL of digestion buffer (50 mM TEAB) were added onto the filter and digestion was carried out at 37°C for 6 h. To elute peptides, three stepwise buffers were applied, with 200 μL of each with one more repeat, including 50 mM TEAB, 0.2% formic acid in H2O, and 50% acetonitrile and 0.2% formic acid in H2O. The peptide solutions were pooled, lyophilized, and resuspended in 500LJμL of 0.1% FA. Those samples were diluted a further 10-fold with 0.1% FA.

Desalted peptides were adjusted to a protein concentration of 100 ng/uL with 0.1% formic acid (FA) in preparation for MS analysis. 150 ng of Digested peptides were loaded into autosampler vials and analyzed directly using a NanoElute liquid chromatography system (Bruker, Germany) coupled with a timsTOF SCP mass spectrometer (Bruker, Germany). Peptides were separated on a 75 μm i.d. × 15 cm separation column packed with 1.9 μm C18 beads (Bruker, Germany) over a 90-minute elution gradient. Buffer A was 0.1% FA in water and buffer B was 0.1% FA in acetonitrile. Instrument control and data acquisition were performed using Compass Hystar (version 6.0) with the timsTOF SCP operating in parallel accumulation serial fragmentation (PASEF) mode under the following settings: mass range 100-1700 m/z, 1/k/0 Start 0.7 V s cm-2 End 1.3 V s cm-2; ramp accumulation times were 166 ms; capillary voltage was 4500 V, dry gas 8.0 L min-1 and dry temp 200°C. The PASEF settings were: 5 MS/MS scans (total cycle time, 1.03 s); charge range 0–5; active exclusion for 0.2 min; scheduling target intensity 20,000; intensity threshold 500; collision-induced dissociation energy 10 eV. The MS/MS data was processed via FragPipe v 20.0 using the SwissProt database with reverse decoys and contaminants added. Oxidation of Met was fixed while carbamidomethylation of Cys was variable with all protein identifications filtered by a 1% false discovery rate (FDR). Identified proteins were sorted into downregulated and upregulated, using a ±1.5 fold change cutoff. Pathway enrichment on each set of up- and down-regulated proteins was analyzed via ShinyGO 0.77 (Ge SX, Jung D & Yao R, <u>Bioinformatics 36:2628–2629, 2020.</u>) to generate respective pathways with an FDR cutoff of 5%.

### 5.14 Statistical Analysis

For all analysis where two groups were compared, t-tests were performed and statistical significance stated where p < 0.05 and trends stated where p < 0.10. When two or more groups were analyzed and compared, a one-way analysis of variance (ANOVA), or two-way ANOVA was performed with Tukey post-hoc analysis comparing all groups, using an appropriately conservative correction for multiple comparisons. Statistical analyses were carried out using GraphPad Prism 9.

## Abbreviations

AD: Alzheimer’s disease
FLIM: Fluorescence Lifetime Imaging Microscopy
DAM: disease-associated microglia
E2: 17β-estradiol
LPL: Lipoprotein lipase
MDMi: monocyte-derived microglia-like cells
NFT: neurofibrillary tangles
LD: lipid droplets
PET: positron emission tomography
Aβ: amyloid beta
TLR: toll-like receptor
ApoE: apolipoprotein E
NADH: nicotinamide adenine dinucleotide
FAD: flavin adenine dinucleotide
FLIRR: Fluorescence Lifetime Imaging Redox Ratio
OXPHOS: oxidative phosphorylation
ATP: adenosine triphosphate
TCA: tricarboxylic acid
CD11b: cluster of differentiation molecule 11b
FBS: fetal bovine serum
ESR1: estrogen receptor 1
ERα: estrogen receptor alpha
ESR2: estrogen receptor 2
ERβ: estrogen receptor beta
PPT: 4,4’,4’’-(4-Propyl-[1*H*]-pyrazole-1,3,5-triyl)*tris*phenol
DPN: diarylpropionitrile
Angptl4: angiopoietin-like 4
F1,6BP: fructose-1,6-bisphosphate
G3P: glyceraldehyde 3-phosphate
OAA: oxaloacetate
PEP: phosphoenolpyruvate
2-OGM: 2- oxoglutaramate
ADP: adenosine diphosphate
iPSCs: induced pluripotent stem cells
PBMC: peripheral blood mononuclear cell
CXCL31: chemokine ligand 31
CD200: cluster of differentiation 200
Trem2: triggering receptor expressed on myeloid cells 2
Iba1: ionized calcium-binding adapter molecule 1
IL-1β: interleukin-1beta
HRT: hormone replacement therapy
FSH: follicle stimulating hormone
GPER: G-protein coupled estrogen receptor
DMEM: Dulbecco’s Modified Eagle Medium
DMSO: dimethyl sulfoxide
MACs: magnetic cell separation
qPCR: quantitative polymerase chain reaction
KH: KRP-heparin solution
LH: lysis-heparin

## Author Contributions

**Nicholas Cleland:** Investigation, Writing – Original Draft, Visualization, Methodology, Validation, Formal analysis, Review & Editing. **Garrett Potter:** Methodology, Investigation, Resources, Analysis. **Daphne Quang:** Methodology, Visualization, Investigation. **Courtney Buck:** Investigation, Methodology. **Dean Oldham:** Resources, Methodology. **Mikaela Neal:** Methodology. **Christy Niemeyer:** Methodology, Resources **Evgenia Dobrinskikh:** Investigation, Methodology, Software, Visualization, Supervision, Formal analysis. **Kimberley Bruce:** Conceptualization, Funding acquisition, Project Administration, Supervision, Data curation, Writing – Original draft, Review & Editing.

## Acknowledgements

We would like to thank Dr. Steven Sheridan for his advice and guidance regarding MDMi cultures. We would also like to thank the University of Colorado School of Medicine Metabolomics Core Facility as well as the Genomics and Microarray Shared Resource which are supported by the Cancer Center Support Grant (P30CA046934). Fluorescent Lifetime Imaging experiments were performed in the University of Colorado Anschutz Medical Campus Advance Light Microscopy Core supported in part by Rocky Mountain Neurological Disorders Core Grant Number P30NS048154.

---

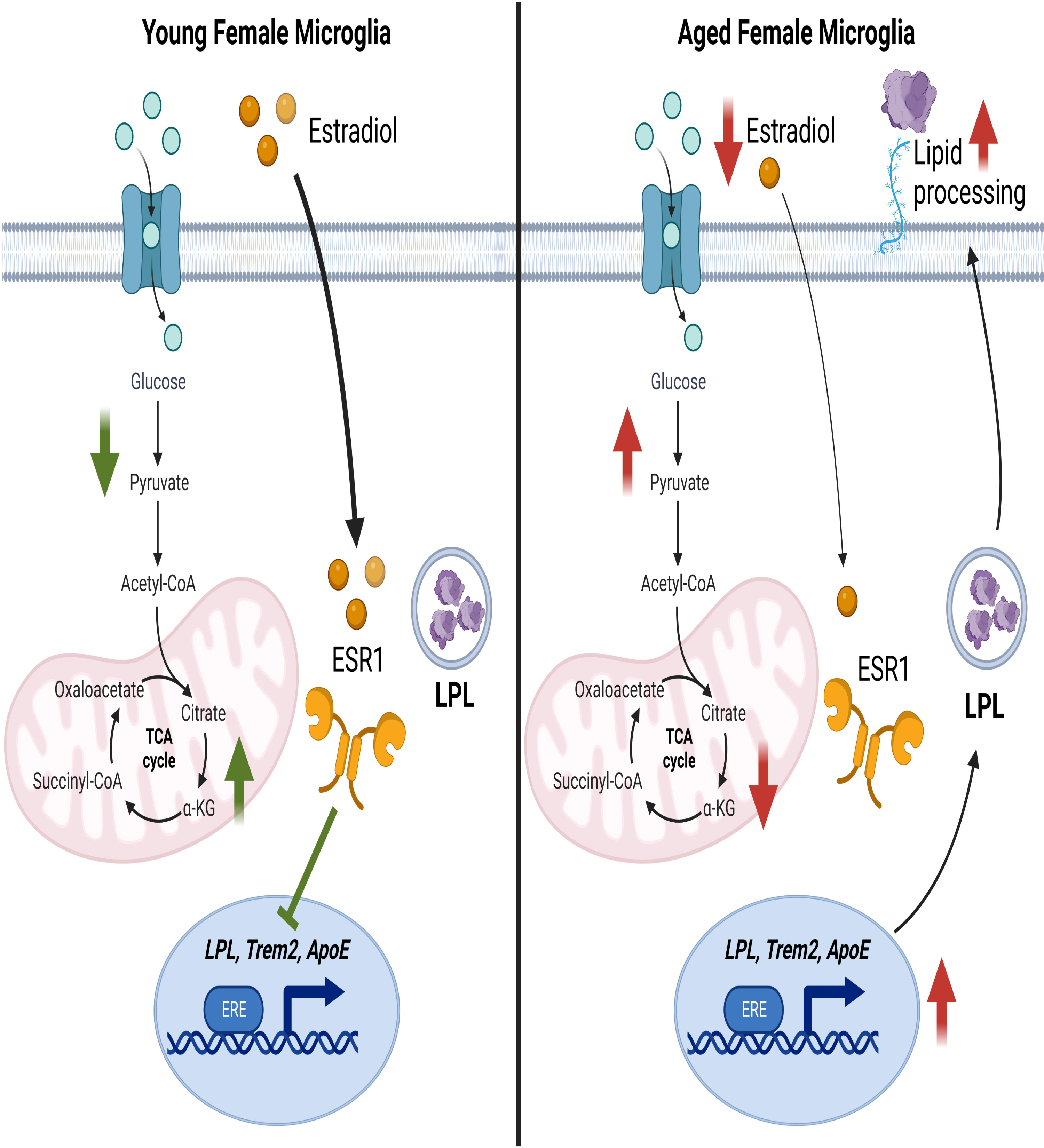

## References

Abeysinghe, A.A.D.T., Deshapriya, R.D.U.S., Udawatte, C., 2020. Alzheimer’s disease; a review of the pathophysiological basis and therapeutic interventions. Life Sciences. 256, 117996.

Abud, E.M., et al., 2017. iPSC-Derived Human Microglia-like Cells to Study Neurological Diseases. Neuron. 94, 278–293.e9.

Angst, G., Tang, X., Wang, C., 2023. Functional Analysis of a Novel Immortalized Murine Microglia Cell Line in 3D Spheroid Model. Neurochem Res.

Baik, S.H., et al., 2019. A Breakdown in Metabolic Reprogramming Causes Microglia Dysfunction in Alzheimer’s Disease. Cell Metab. 30, 493–507 e6.

Baker, A.E., Brautigam, V.M., Watters, J.J., 2004. Estrogen modulates microglial inflammatory mediator production via interactions with estrogen receptor beta. Endocrinology. 145, 5021–32.

Berezin, M.Y., Achilefu, S., 2010. Fluorescence lifetime measurements and biological imaging. Chemical reviews. 110, 2641–2684.

Bruce-Keller, A.J., et al., 2000. Antiinflammatory effects of estrogen on microglial activation. Endocrinology. 141, 3646–56.

Burant, C.F., et al., 1992. Fructose transporter in human spermatozoa and small intestine is GLUT5. J Biol Chem. 267, 14523–6.

Butovsky, O., Weiner, H.L., 2018. Microglial signatures and their role in health and disease. Nat Rev Neurosci. 19, 622–635.

Calsolaro, V., Edison, P., 2016. Neuroinflammation in Alzheimer’s disease: Current evidence and future directions. Alzheimers Dement. 12, 719–32.

Cantuti-Castelvetri, I., et al., 2007. Effects of gender on nigral gene expression and parkinson disease. Neurobiol Dis. 26, 606–14.

Cao, R., et al., 2019. Single-cell redox states analyzed by fluorescence lifetime metrics and tryptophan FRET interaction with NAD(P)H. Cytometry A. 95, 110–121.

Chen, H., et al., 2023. The immunometabolic reprogramming of microglia in Alzheimer’s disease. Neurochem Int. 171, 105614.

Cleland, N.R.W., et al., 2021. Altered substrate metabolism in neurodegenerative disease: new insights from metabolic imaging. J Neuroinflammation. 18, 248.

Coales, I., et al., 2022. Alzheimer’s disease-related transcriptional sex differences in myeloid cells. Journal of Neuroinflammation. 19, 247.

de Luna, N., et al., 2023. Neuroinflammation-Related Proteins NOD2 and Spp1 Are Abnormally Upregulated in Amyotrophic Lateral Sclerosis. Neurol Neuroimmunol Neuroinflamm. 10.

De Schepper, S., et al., 2023. Perivascular cells induce microglial phagocytic states and synaptic engulfment via SPP1 in mouse models of Alzheimer’s disease. Nat Neurosci. 26, 406–415.

De Sousa, R.A.L., 2022. Reactive gliosis in Alzheimer’s disease: a crucial role for cognitive impairment and memory loss. Metab Brain Dis. 37, 851–857.

Demarest, T.G., et al., 2020. Biological sex and DNA repair deficiency drive Alzheimer’s disease via systemic metabolic remodeling and brain mitochondrial dysfunction. Acta Neuropathol. 140, 25–47.

Digman, M.A., et al., 2008. The phasor approach to fluorescence lifetime imaging analysis. Biophysical journal. 94, L14–L16.

Dimayuga, F.O., et al., 2005. Estrogen and brain inflammation: effects on microglial expression of MHC, costimulatory molecules and cytokines. J Neuroimmunol. 161, 123–36.

Dobin, A., et al., 2013. STAR: ultrafast universal RNA-seq aligner. Bioinformatics. 29, 15–21.

Dobrinskikh, E., et al., 2019. Heterogeneous pulmonary response after tracheal occlusion: clues to fetal lung growth. Journal of Surgical Research. 239, 242–252.

Doolittle, M.H., et al., 1990. The response of lipoprotein lipase to feeding and fasting. Evidence for posttranslational regulation. J Biol Chem. 265, 4570–7.

Duncan, K.A., Saldanha, C.J., 2011. Neuroinflammation induces glial aromatase expression in the uninjured songbird brain. J Neuroinflammation. 8, 81.

Erkkilä, M.T., et al., 2020. Macroscopic fluorescence-lifetime imaging of NADH and protoporphyrin IX improves the detection and grading of 5-aminolevulinic acid-stained brain tumors. Scientific Reports. 10, 20492.

Fairley, L.H., Wong, J.H., Barron, A.M., 2021. Mitochondrial Regulation of Microglial Immunometabolism in Alzheimer’s Disease. Front Immunol. 12, 624538.

Fumagalli, M., et al., 2018. How to reprogram microglia toward beneficial functions. Glia. 66, 2531–2549.

Gold, S.M., et al., 2009. Estrogen treatment decreases matrix metalloproteinase (MMP)-9 in autoimmune demyelinating disease through estrogen receptor alpha (ERalpha). Lab Invest. 89, 1076–83.

Goyal, M.S., et al., 2014. Aerobic glycolysis in the human brain is associated with development and neotenous gene expression. Cell Metab. 19, 49–57.

Goyal, M.S., et al., 2020. Spatiotemporal relationship between subthreshold amyloid accumulation and aerobic glycolysis in the human brain. Neurobiol Aging. 96, 165–175.

Grady, D., et al., 2002. Effect of postmenopausal hormone therapy on cognitive function: the Heart and Estrogen/progestin Replacement Study. Am J Med. 113, 543–8.

Grubman, A., et al., 2021. Transcriptional signature in microglia associated with Abeta plaque phagocytosis. Nat Commun. 12, 3015.

Guan, J., et al., 2017. GPER Agonist G1 Attenuates Neuroinflammation and Dopaminergic Neurodegeneration in Parkinson Disease. Neuroimmunomodulation. 24, 60–66.

Guillot-Sestier, M.V., et al., 2021. Microglial metabolism is a pivotal factor in sexual dimorphism in Alzheimer’s disease. Commun Biol. 4, 711.

Habermehl, T.L., et al., 2022. Aging-associated changes in motor function are ovarian somatic tissue-dependent, but germ cell and estradiol independent in post-reproductive female mice exposed to young ovarian tissue. GeroScience. 44, 2157–2169.

Hammond, T.R., et al., 2019. Single-Cell RNA Sequencing of Microglia throughout the Mouse Lifespan and in the Injured Brain Reveals Complex Cell-State Changes. Immunity. 50, 253–271.e6.

Hamosh, M., Hamosh, P., 1975. The effect of estrogen on the lipoprotein lipase activity of rat adipose tissue. The Journal of Clinical Investigation. 55, 1132–1135.

Han, J., et al., 2023. Label-Free Characterization of Atherosclerotic Plaques Via High-Resolution Multispectral Fluorescence Lifetime Imaging Microscopy. Arteriosclerosis, Thrombosis, and Vascular Biology. 43, 1295–1307.

Hanamsagar, R., et al., 2017. Generation of a microglial developmental index in mice and in humans reveals a sex difference in maturation and immune reactivity. Glia. 65, 1504–1520.

Hariharan, V.A., et al., 2017. The Enzymology of 2-Hydroxyglutarate, 2-Hydroxyglutaramate and 2- Hydroxysuccinamate and Their Relationship to Oncometabolites. Biology (Basel). 6.

Heneka, M.T., et al., 2015. Neuroinflammation in Alzheimer’s disease. Lancet Neurol. 14, 388–405.

Henn, A., et al., 2009. The suitability of BV2 cells as alternative model system for primary microglia cultures or for animal experiments examining brain inflammation. Altex. 26, 83–94.

Homma, H., et al., 2000. Estrogen Suppresses Transcription of Lipoprotein Lipase Gene: EXISTENCE OF A UNIQUE ESTROGEN RESPONSE ELEMENT ON THE LIPOPROTEIN LIPASE PROMOTER*. Journal of Biological Chemistry. 275, 11404–11411.

Iverius, P.H., Brunzell, J.D., 1988. Relationship between lipoprotein lipase activity and plasma sex steroid level in obese women. The Journal of Clinical Investigation. 82, 1106–1112.

Johri, A., 2021. Disentangling Mitochondria in Alzheimer’s Disease. International Journal of Molecular Sciences. 22, 11520.

Jung, E.S., et al., 2022. Amyloid-beta activates NLRP3 inflammasomes by affecting microglial immunometabolism through the Syk-AMPK pathway. Aging Cell. 21, e13623.

Kao, Y.C., et al., 2020. Lipids and Alzheimer’s Disease. Int J Mol Sci. 21.

Kim, E.J., et al., 2005. Glucose metabolism in early onset versus late onset Alzheimer’s disease: an SPM analysis of 120 patients. Brain. 128, 1790–1801.

Kim, H.J., Kalkhoff, R.K., 1975. Sex steroid influence on triglyceride metabolism. The Journal of Clinical Investigation. 56, 888–896.

Kim, O.Y., et al., 2012. Effects of aging and menopause on serum interleukin-6 levels and peripheral blood mononuclear cell cytokine production in healthy nonobese women. Age (Dordr). 34, 415–25.

Kim, S., et al., 2022. Brain Region-Dependent Alternative Splicing of Alzheimer Disease (AD)-Risk Genes Is Associated With Neuropathological Features in AD. Int Neurourol J. 26, S126–136.

Kodama, L., Gan, L., 2019. Do Microglial Sex Differences Contribute to Sex Differences in Neurodegenerative Diseases? Trends Mol Med. 25, 741–749.

Kooner, J.S., et al., 2008. Genome-wide scan identifies variation in MLXIPL associated with plasma triglycerides. Nature Genetics. 40, 149–151.

Lee, D.H., et al., 2017. Age-dependent alterations in serum cytokines, peripheral blood mononuclear cell cytokine production, natural killer cell activity, and prostaglandin F(2α). Immunol Res. 65, 1009–1016.

Liao, Y., Smyth, G.K., Shi, W., 2014. featureCounts: an efficient general purpose program for assigning sequence reads to genomic features. Bioinformatics. 30, 923–30.

Liu, K.Y., Howard, R., 2021. Can we learn lessons from the FDA’s approval of aducanumab? Nat Rev Neurol. 17, 715–722.

Lloyd, A.F., et al., 2019. Central nervous system regeneration is driven by microglia necroptosis and repopulation. Nat Neurosci. 22, 1046–1052.

Loiola, R.A., et al., 2019. Estrogen Promotes Pro-resolving Microglial Behavior and Phagocytic Cell Clearance Through the Actions of Annexin A1. Frontiers in Endocrinology. 10.

Loram, L.C., et al., 2012. Sex and estradiol influence glial pro-inflammatory responses to lipopolysaccharide in rats. Psychoneuroendocrinology. 37, 1688–99.

Love, M.I., Huber, W., Anders, S., 2014. Moderated estimation of fold change and dispersion for RNA- seq data with DESeq2. Genome Biol. 15, 550.

Loving, B.A., et al., 2021. Lipoprotein Lipase Regulates Microglial Lipid Droplet Accumulation. Cells. 10.

Low, H., Hoang, A., Sviridov, D., 2012. Cholesterol efflux assay. JoVE (Journal of Visualized Experiments). e3810.

Lukina, M., et al., 2021. Label-Free Macroscopic Fluorescence Lifetime Imaging of Brain Tumors. Frontiers in Oncology. 11.

Lynch, M.A., 2020. Can the emerging field of immunometabolism provide insights into neuroinflammation? Progress in Neurobiology. 184, 101719.

Lynch, M.A., 2022. Exploring Sex-Related Differences in Microglia May Be a Game-Changer in Precision Medicine. Frontiers in Aging Neuroscience. 14.

Manczak, M., et al., 2004. Differential expression of oxidative phosphorylation genes in patients with Alzheimer’s disease: implications for early mitochondrial dysfunction and oxidative damage. Neuromolecular Med. 5, 147–62.

Manly, J.J., Glymour, M.M., 2021. What the Aducanumab Approval Reveals About Alzheimer Disease Research. JAMA Neurol. 78, 1305–1306.

Marschallinger, J., et al., 2020. Lipid-droplet-accumulating microglia represent a dysfunctional and proinflammatory state in the aging brain. Nat Neurosci. 23, 194–208.

Marwan, A.I., et al., 2019. Unique heterogeneous topological pattern of the metabolic landscape in rabbit fetal lungs following tracheal occlusion. Fetal diagnosis and therapy. 45, 145–154.

Masters, C.L., et al., 1985. Amyloid plaque core protein in Alzheimer disease and Down syndrome. Proc Natl Acad Sci U S A. 82, 4245–9.

McCurdy, C.E., et al., 2016. Maternal obesity reduces oxidative capacity in fetal skeletal muscle of Japanese macaques. JCI insight. 1.

McQuade, A., et al., 2018. Development and validation of a simplified method to generate human microglia from pluripotent stem cells. Mol Neurodegener. 13, 67.

Mei, Y., et al., 2009. FOXO3a-dependent regulation of Pink1 (Park6) mediates survival signaling in response to cytokine deprivation. Proceedings of the National Academy of Sciences. 106, 5153–5158.

Melamud, E., Vastag, L., Rabinowitz, J.D., 2010. Metabolomic analysis and visualization engine for LC− MS data. Analytical chemistry. 82, 9818–9826.

Minhas, P.S., et al., 2021. Restoring metabolism of myeloid cells reverses cognitive decline in ageing. Nature. 590, 122–128.

Mullins, R., Reiter, D., Kapogiannis, D., 2018. Magnetic resonance spectroscopy reveals abnormalities of glucose metabolism in the Alzheimer’s brain. Ann Clin Transl Neurol. 5, 262–272.

Mulnard, R.A., et al., 2000. Estrogen Replacement Therapy for Treatment of Mild to Moderate Alzheimer DiseaseA Randomized Controlled Trial. JAMA. 283, 1007–1015.

Nelson, L.H., Warden, S., Lenz, K.M., 2017. Sex differences in microglial phagocytosis in the neonatal hippocampus. Brain Behav Immun. 64, 11–22.

Oakley, H., et al., 2006. Intraneuronal beta-amyloid aggregates, neurodegeneration, and neuron loss in transgenic mice with five familial Alzheimer’s disease mutations: potential factors in amyloid plaque formation. J Neurosci. 26, 10129–40.

Ocanas, S.R., et al., 2023. Microglial senescence contributes to female-biased neuroinflammation in the aging mouse hippocampus: implications for Alzheimer’s disease. J Neuroinflammation. 20, 188.

Olah, M., et al., 2020. Single cell RNA sequencing of human microglia uncovers a subset associated with Alzheimer’s disease. Nat Commun. 11, 6129.

Oldham, D., et al., 2022. Using Synthetic ApoC-II Peptides and nAngptl4 Fragments to Measure Lipoprotein Lipase Activity in Radiometric and Fluorescent Assays. Front Cardiovasc Med. 9, 926631.

Osborne, O.M., et al., 2023. Anti-amyloid: An antibody to cure Alzheimer’s or an attitude. iScience. 26, 107461.

Paik, J.K., et al., 2012. Circulating and PBMC Lp-PLA2 associate differently with oxidative stress and subclinical inflammation in nonobese women (menopausal status). PLoS One. 7, e29675.

Park, J.C., et al., 2023. Sex differences in the progression of glucose metabolism dysfunction in Alzheimer’s disease. Exp Mol Med. 55, 1023–1032.

Pedersen, S.B., et al., 1991. Nuclear estradiol binding in rat adipocytes. Regional variations and regulatory influences of hormones. Biochimica et Biophysica Acta (BBA) - Molecular Cell Research. 1093, 80–86.

Pedersen, S.B., et al., 1992. Effects of in vivo estrogen treatment on adipose tissue metabolism and nuclear estrogen receptor binding in isolated rat adipocytes. Molecular and Cellular Endocrinology. 85, 13–19.

Peinado-Onsurbe, J., et al., 2008. Effects of Sex Steroids on Hepatic and Lipoprotein Lipase Activity and mRNA in the Rat. Hormone Research. 40, 184–188.

Porsteinsson, A.P., et al., 2021. Diagnosis of Early Alzheimer’s Disease: Clinical Practice in 2021. J Prev Alzheimers Dis. 8, 371–386.

Preeti, K., Sood, A., Fernandes, V., 2022. Metabolic Regulation of Glia and Their Neuroinflammatory Role in Alzheimer’s Disease. Cell Mol Neurobiol. 42, 2527–2551.

Qu, Y., et al., 2022. Estrogen Up-Regulates Iron Transporters and Iron Storage Protein Through Hypoxia Inducible Factor 1 Alpha Activation Mediated by Estrogen Receptor β and G Protein Estrogen Receptor in BV2 Microglia Cells. Neurochemical Research. 47, 3659–3669.

Rahman, A., et al., 2019. Sex and Gender Driven Modifiers of Alzheimer’s: The Role for Estrogenic Control Across Age, Race, Medical, and Lifestyle Risks. Frontiers in Aging Neuroscience. 11.

Rajan, K.B., et al., 2021. Population estimate of people with clinical Alzheimer’s disease and mild cognitive impairment in the United States (2020-2060). Alzheimers Dement. 17, 1966–1975.

Ramanan, V.K., Day, G.S., 2023. Anti-amyloid therapies for Alzheimer disease: finally, good news for patients. Mol Neurodegener. 18, 42.

Ranjit, S., et al., 2017. Measuring the effect of a Western diet on liver tissue architecture by FLIM autofluorescence and harmonic generation microscopy. Biomed Opt Express. 8, 3143–3154.

Riedel, B.C., Thompson, P.M., Brinton, R.D., 2016. Age, APOE and sex: Triad of risk of Alzheimer’s disease. J Steroid Biochem Mol Biol. 160, 134–47.

Sagar, M.A.K., et al., 2020. Microglia activation visualization via fluorescence lifetime imaging microscopy of intrinsically fluorescent metabolic cofactors. Neurophotonics. 7, 035003.

Saijo, K., et al., 2011. An ADIOL-ERβ-CtBP transrepression pathway negatively regulates microglia-mediated inflammation. Cell. 145, 584–95.

Sala Frigerio, C., et al., 2019. The Major Risk Factors for Alzheimer’s Disease: Age, Sex, and Genes Modulate the Microglia Response to Aβ Plaques. Cell Rep. 27, 1293–1306.e6.

Saleh, R.N.M., et al., 2023. Hormone replacement therapy is associated with improved cognition and larger brain volumes in at-risk APOE4 women: results from the European Prevention of Alzheimer’s Disease (EPAD) cohort. Alzheimers Res Ther. 15, 10.

Schwarz, J.M., Sholar, P.W., Bilbo, S.D., 2012. Sex differences in microglial colonization of the developing rat brain. J Neurochem. 120, 948–63.

Sellgren, C.M., et al., 2017. Patient-specific models of microglia-mediated engulfment of synapses and neural progenitors. Mol Psychiatry. 22, 170–177.

Sepehr, E., et al., 2012. Pharmacokinetics of the estrogen receptor subtype-selective ligands, PPT and DPN: quantification using UPLC-ES/MS/MS. J Pharm Biomed Anal. 71, 119–26.

Serrano-Pozo, A., et al., 2011. Neuropathological alterations in Alzheimer disease. Cold Spring Harb Perspect Med. 1, a006189.

Shang, R., Rodrigues, B., 2021. Lipoprotein Lipase and Its Delivery of Fatty Acids to the Heart. Biomolecules. 11.

Sheridan, S.D., Horng, J.E., Perlis, R.H., 2022. Patient-Derived In Vitro Models of Microglial Function and Synaptic Engulfment in Schizophrenia. Biol Psychiatry. 92, 470–479.

Shippy, D.C., Ulland, T.K., 2020. Microglial Immunometabolism in Alzheimer’s Disease. Front Cell Neurosci. 14, 563446.

Sierksma, A., et al., 2020. Novel Alzheimer risk genes determine the microglia response to amyloid-β but not to TAU pathology. EMBO Mol Med. 12, e10606.

Slowik, A., et al., 2018. Impact of steroid hormones E2 and P on the NLRP3/ASC/Casp1 axis in primary mouse astroglia and BV-2 cells after in vitro hypoxia. J Steroid Biochem Mol Biol. 183, 18–26.

Tatibouët, A., et al., 2000. Synthesis and evaluation of fructose analogues as inhibitors of the D-fructose transporter GLUT5. Bioorg Med Chem. 8, 1825–33.

Thion, M.S., et al., 2018. Microbiome Influences Prenatal and Adult Microglia in a Sex-Specific Manner. Cell. 172, 500–516.e16.

Timmerman, R., Burm, S.M., Bajramovic, J.J., 2018. An Overview of in vitro Methods to Study Microglia. Front Cell Neurosci. 12, 242.

Vaishnavi, S.N., et al., 2010. Regional aerobic glycolysis in the human brain. Proc Natl Acad Sci U S A. 107, 17757–62.

van Dyck, C.H., et al., 2023. Lecanemab in Early Alzheimer’s Disease. N Engl J Med. 388, 9–21.

Vegeto, E., et al., 2000. Estrogen blocks inducible nitric oxide synthase accumulation in LPS-activated microglia cells. Exp Gerontol. 35, 1309–16.

Vegeto, E., et al., 2001. Estrogen prevents the lipopolysaccharide-induced inflammatory response in microglia. J Neurosci. 21, 1809–18.

Villa, A., et al., 2018. Sex-Specific Features of Microglia from Adult Mice. Cell Reports. 23, 3501–3511.

Vinogradova, Y., Coupland, C., Hippisley-Cox, J., 2019. Use of hormone replacement therapy and risk of venous thromboembolism: nested case-control studies using the QResearch and CPRD databases. Bmj. 364, k4810.

Wallrabe, H., et al., 2018. Segmented cell analyses to measure redox states of autofluorescent NAD (P) H, FAD & Trp in cancer cells by FLIM. Scientific reports. 8, 1–11.

Wang, J., et al., 2021. Estrogen Attenuates Traumatic Brain Injury by Inhibiting the Activation of Microglia and Astrocyte-Mediated Neuroinflammatory Responses. Mol Neurobiol. 58, 1052–1061.

Wang, S., et al., 2012. Mitochondrial fission proteins in peripheral blood lymphocytes are potential biomarkers for Alzheimer’s disease. European Journal of Neurology. 19, 1015–1022.

Wang, X., et al., 2014. Oxidative stress and mitochondrial dysfunction in Alzheimer’s disease. Biochim Biophys Acta. 1842, 1240–7.

Wood, J.G., et al., 1986. Neurofibrillary tangles of Alzheimer disease share antigenic determinants with the axonal microtubule-associated protein tau (tau). Proc Natl Acad Sci U S A. 83, 4040–3.

Wu, M., et al., 2020. Postmenopausal hormone therapy and Alzheimer’s disease, dementia, and Parkinson’s disease: A systematic review and time-response meta-analysis. Pharmacol Res. 155, 104693.

Wu, W.F., et al., 2013. Targeting estrogen receptor β in microglia and T cells to treat experimental autoimmune encephalomyelitis. Proc Natl Acad Sci U S A. 110, 3543–8.

Xia, J., Wishart, D.S., 2010a. MSEA: a web-based tool to identify biologically meaningful patterns in quantitative metabolomic data. Nucleic acids research. 38, W71–W77.

Xia, J., Wishart, D.S., 2010b. MetPA: a web-based metabolomics tool for pathway analysis and visualization. Bioinformatics. 26, 2342–2344.

Xiong, J., et al., 2022. FSH blockade improves cognition in mice with Alzheimer’s disease. Nature. 603, 470–476.

Yanguas-Casás, N., et al., 2018. Sex differences in the phagocytic and migratory activity of microglia and their impairment by palmitic acid. Glia. 66, 522–537.

Yun, J., et al., 2018. Estrogen deficiency exacerbates Aβ-induced memory impairment through enhancement of neuroinflammation, amyloidogenesis and NF-ĸB activation in ovariectomized mice. Brain Behav Immun. 73, 282–293.

Zhang, G.Q., et al., 2021. Menopausal hormone therapy and women’s health: An umbrella review. PLoS Med. 18, e1003731.

Zhang, Y., et al., 2014. An RNA-sequencing transcriptome and splicing database of glia, neurons, and vascular cells of the cerebral cortex. J Neurosci. 34, 11929–47.

Zhang, Y., et al., 2016. Purification and Characterization of Progenitor and Mature Human Astrocytes Reveals Transcriptional and Functional Differences with Mouse. Neuron. 89, 37–53.

Zhao, L., et al., 2016. Sex differences in metabolic aging of the brain: insights into female susceptibility to Alzheimer’s disease. Neurobiol Aging. 42, 69–79.

