## Supplemental Figures for "Altered Metabolism and DAM-signatures in Female Brains and Microglia with Aging": Supplemental Table 6.docx

| **Enrichment FDR** | **nGenes** | **Pathway Genes** | **Fold Enrichment** | **Pathway** | **Genes** |
| --- | --- | --- | --- | --- | --- |
| 0.002307905 | 5 | 44 | 12.33549784 | Aminoacyl-tRNA biosynthesis | KARS1 AARS1 GARS1 WARS1 FARSA |
| 0.037909302 | 3 | 31 | 10.50506912 | Galactose metabolism | PGM1 HK1 UGP2 |
| 0.045867682 | 3 | 35 | 9.304489796 | Starch and sucrose metabolism | PGM1 HK1 UGP2 |
| 0.018553972 | 4 | 49 | 8.861418853 | Amino sugar and nucleotide sugar metabolism | PGM1 CHIT1 HK1 UGP2 |
| 0.018593127 | 4 | 50 | 8.684190476 | Cholesterol metabolism | LRP1 APOC1 NCEH1 LPL |
| 0.000488773 | 9 | 134 | 7.290831557 | Ribosome | RPLP0 RPS16 RPL28 RPS13 RPS15A RPS8 RPS3A RPS14 RPL23A |
| 0.042623996 | 4 | 67 | 6.480739161 | Glycolysis / Gluconeogenesis | PGM1 DLD HK1 FBP1 |
| 0.000861168 | 9 | 151 | 6.470009461 | Phagosome | CD209 TCIRG1 NCF2 ATP6V1G1 DYNC1LI1 ITGA5 ITGAM CALR SEC22B |
| 0.006224136 | 7 | 134 | 5.670646766 | Oxidative phosphorylation | ATP5F1D TCIRG1 ATP5PB ATP6V1G1 ATP5ME COX5A MT-CO2 |
| 0.035956446 | 5 | 102 | 5.321195145 | Amoebiasis | VCL SERPINB3 GNAS LAMB1 ITGAM |
| 0.045867682 | 5 | 115 | 4.719668737 | Carbon metabolism | ME2 DLD HK1 FBP1 HIBCH |
| 0.00231306 | 10 | 232 | 4.678981938 | Coronavirus disease | RPLP0 RPS16 RPL28 RPS13 STAT1 RPS15A RPS8 RPS3A RPS14 RPL23A |
| 0.018553972 | 7 | 169 | 4.496252465 | Protein processing in endoplasmic reticulum | CAPN1 NSFL1C RPN2 RAD23B DAD1 CKAP4 CALR |
| 0.002174355 | 11 | 266 | 4.489008235 | Parkinson disease | GNAS ATP5F1D PSMC1 PPIF ATP5PB TXN SOD1 PSMD2 CALML5 COX5A MT-CO2 |
| 0.006224136 | 9 | 223 | 4.381037796 | Chemical carcinogenesis | MGST2 ATP5F1D PPIF ATP5PB NCF2 SOD1 ACP1 COX5A MT-CO2 |
| 0.003928816 | 10 | 252 | 4.307634165 | Endocytosis | VPS35 RAB10 ARPC3 VPS29 SNX3 WIPF1 ARF3 CLTB IGF2R CAPZA2 |
| 0.000488773 | 15 | 383 | 4.251398732 | Alzheimer disease | CAPN1 HSD17B10 ATP5F1D PSMC1 PPIF RTN4 ATP5PB IDE LRP1 ADAM10 PSMD2 LPL CALML5 COX5A MT-CO2 |
| 0.018553972 | 9 | 272 | 3.591806723 | Prion disease | ATP5F1D PSMC1 PPIF ATP5PB NCF2 SOD1 PSMD2 COX5A MT-CO2 |
| 0.03317122 | 9 | 306 | 3.192717087 | Huntington disease | ATP5F1D PSMC1 PPIF ATP5PB SOD1 PSMD2 CLTB COX5A MT-CO2 |
| 0.008709315 | 28 | 1527 | 1.990482427 | Metabolic pathways | ACSL4 HSD17B10 PGM1 HAL GSTP1 MGST2 DLD ATP5F1D PLD3 PLOD3 TCIRG1 ATP5PB PRDX6 RPN2 DAD1 CHIT1 ATP6V1G1 IMPA2 ACP1 HK1 FBP1 CRYL1 ATP5ME UGP2 COX5A HIBCH MT-CO2 NME1 |
