## Supplementary figures and images for "Altered Metabolism and DAM-signatures in Female Brains and Microglia with Aging"

### Supplemental 2.jpeg

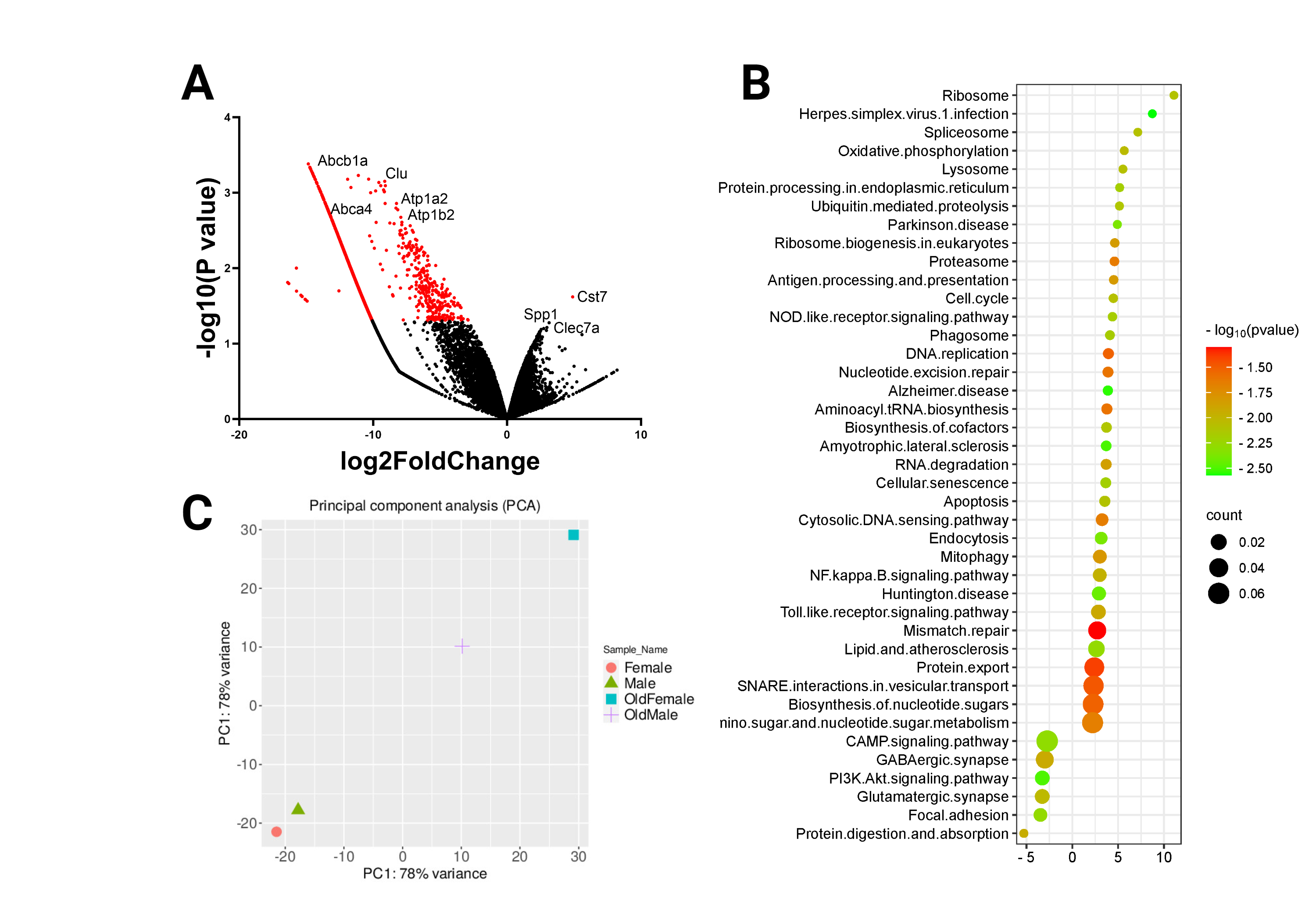

### Supplemental Figure 1_ FLIM maps .jpg

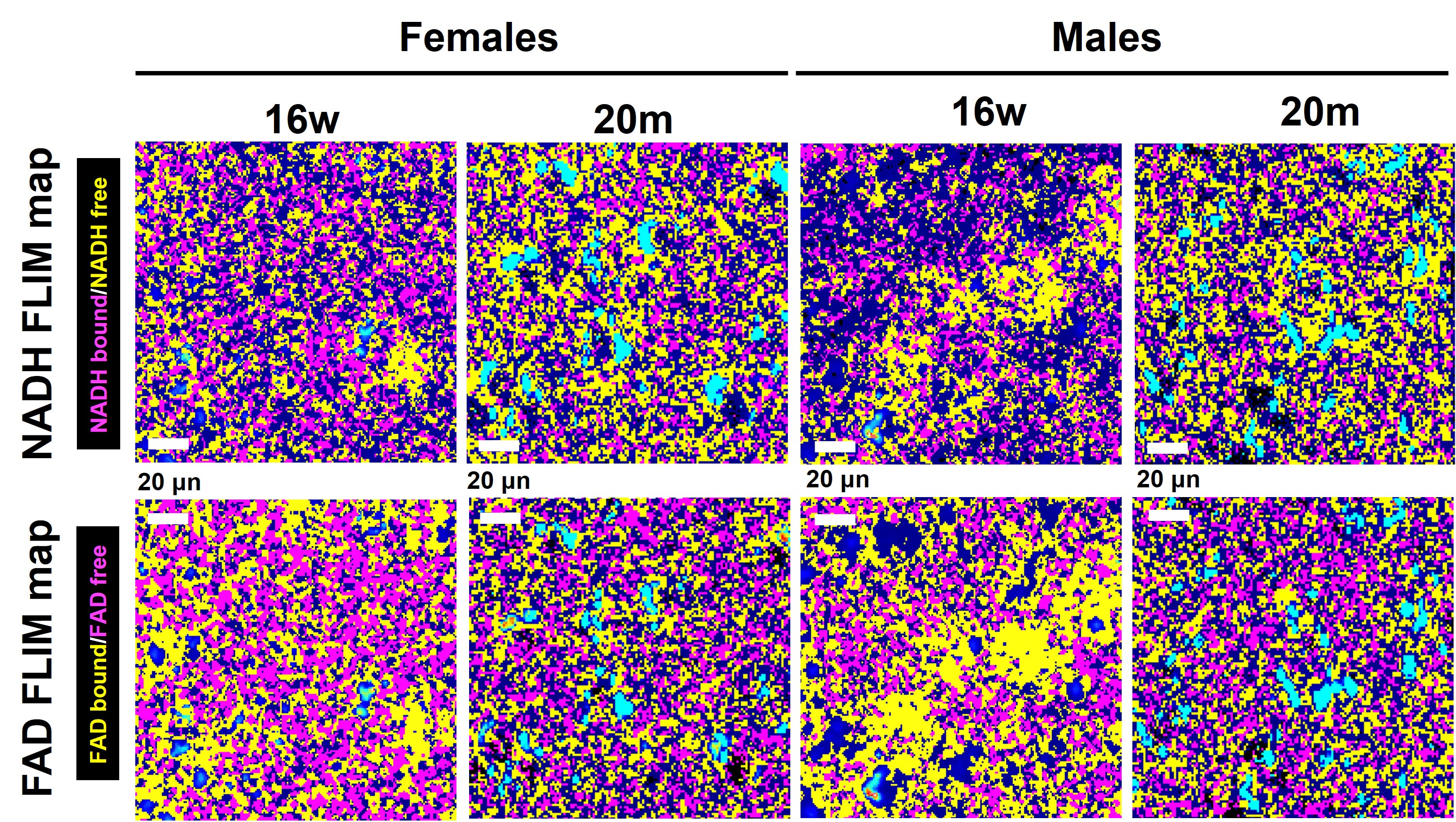

### Supplemental Figure 3.jpg

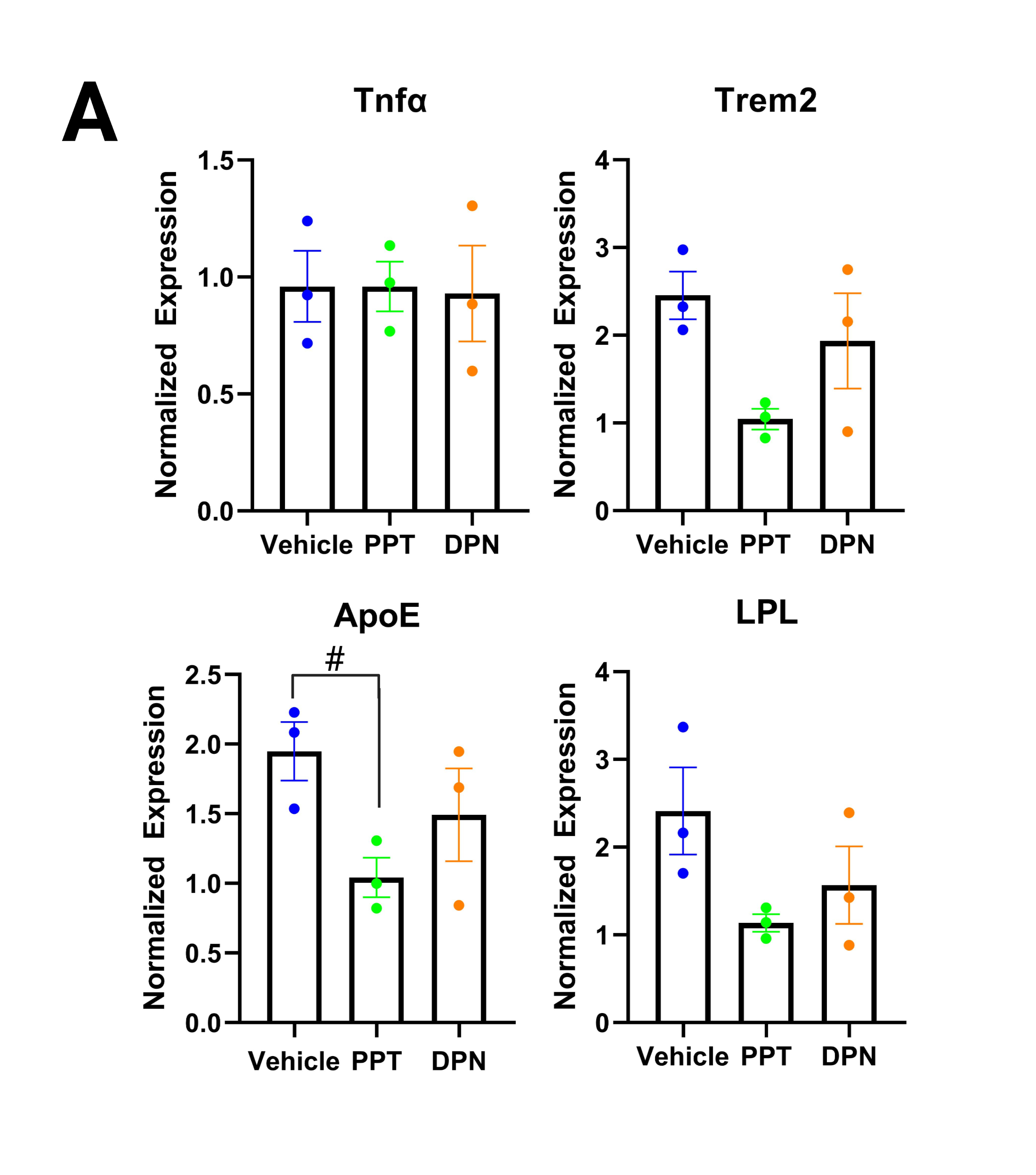

### Supplemental Figure 4.jpg

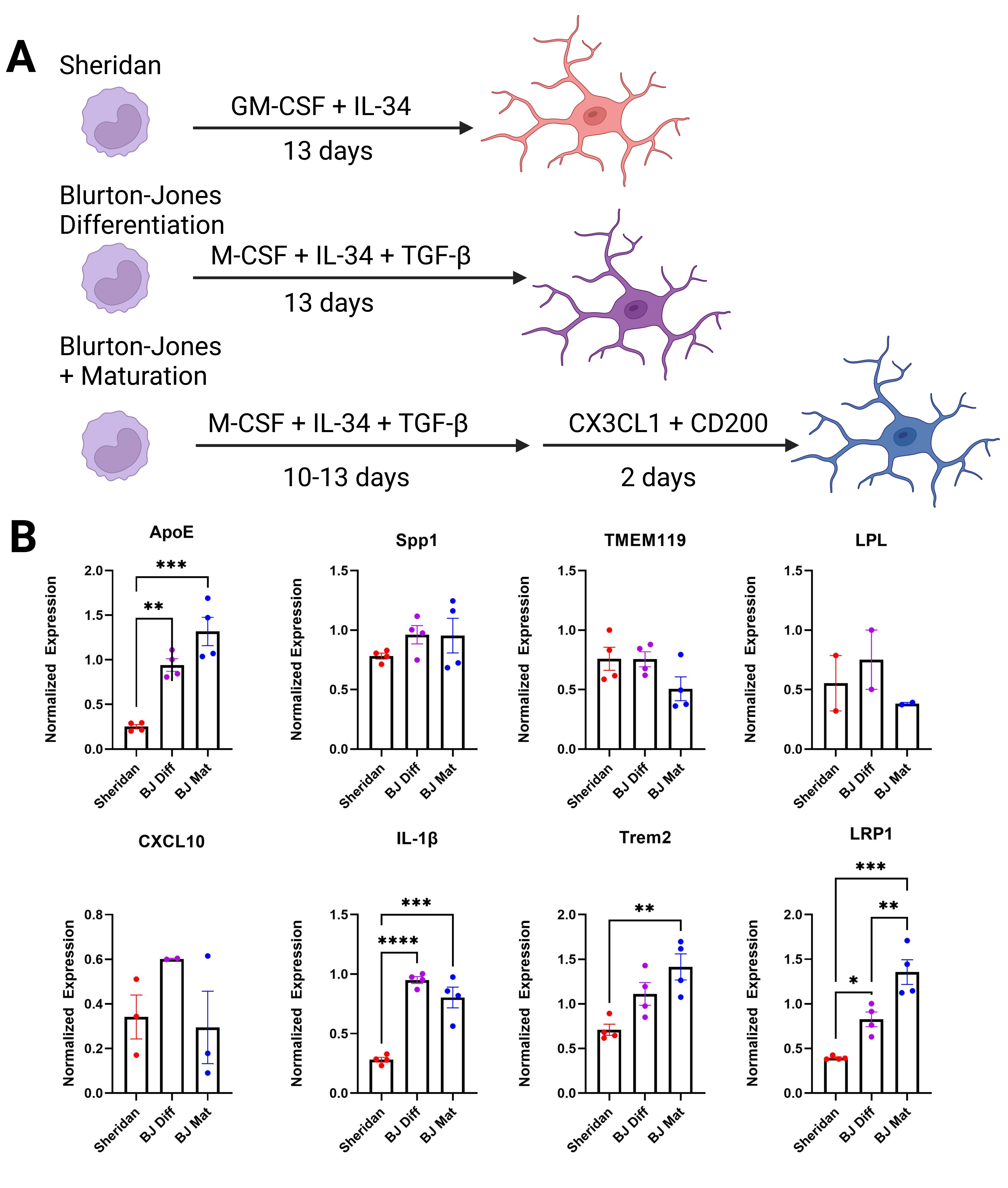

### Supplemental Figure 5_ PBMC Characterization.jpeg

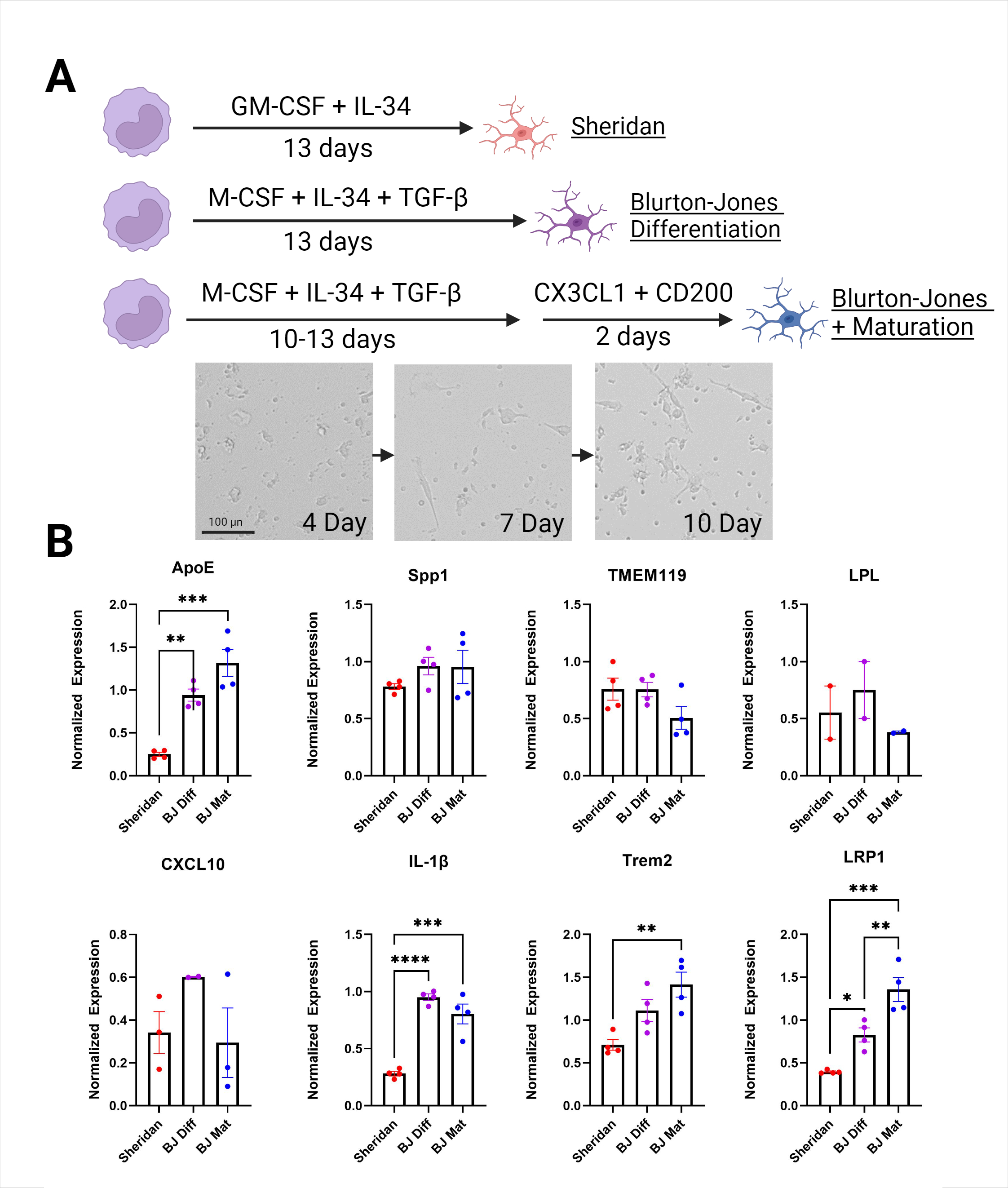

### Supplemental Figure 6.png

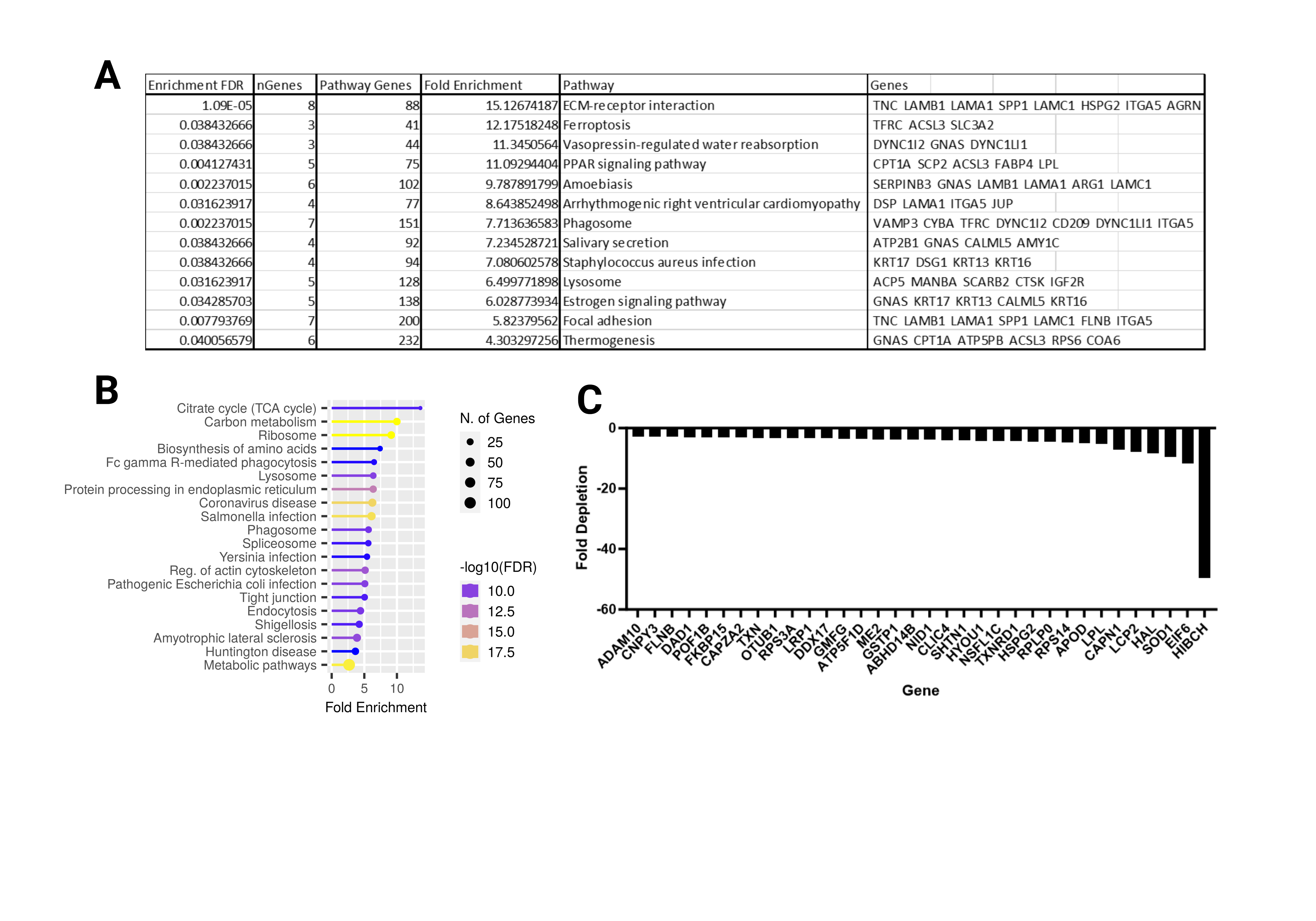
